# Spatial Proteomics Reveals Dual-Compartment Immune Evasion Architecture and Core Lymphomagenic Stromal Niches in Follicular Lymphoma

**DOI:** 10.64898/2026.06.09.731209

**Authors:** Olivier Elemento, Hiranmayi Ravichandran, Wendy Béguelin, David W. Scott, Christian Steidl, Ari M. Melnick

## Abstract

Follicular lymphomas (FL) depend on microenvironmental signals for survival, and the composition of the tumor microenvironment may influence clinical outcome. How it is spatially organized, and how that organization drives immune evasion, has not been established. We applied two complementary 39-marker imaging mass cytometry panels to 156 follicular lymphoma tumor cores, generating a spatial atlas of 4.2 million cells resolved into nine tissue compartments. FL follicles operated as immune-privileged zones: CD8 T cells were largely excluded, and those that did infiltrate were exhausted in a gradient that peaked at the GC core and dissolved at the follicle boundary. A Treg-enriched barrier at that boundary separated the intrafollicular program from an effector-rich interfollicular compartment. Within follicles, CD14-high follicular dendritic cells defined an activated stromal niche that supported tumor B cell proliferation and predicted shorter progression-free and overall survival. Across patients, per-cell CD14 intensity was the strongest predictor of progression, remaining prognostic independently of FLIPI, grade, and stage. VISTA, not PD-L1, dominated checkpoint expression across all myeloid subtypes in both follicular and interfollicular zones, and was further amplified on M2 macrophages in patients carrying EP300 mutations. S100A9+ MDSC-like cells were the strongest myeloid predictor of histologic transformation. Agent-based modeling of this architecture predicted synergistic tumor clearance when anti-VISTA was combined with a CD20-CD3 bispecific antibody. These findings establish that FL uses tissue architecture as an immune evasion mechanism and identify VISTA and CD14-high follicular dendritic cells as candidate therapeutic targets.

## INTRODUCTION

Follicular lymphoma (FL) is the most common indolent non-Hodgkin lymphoma and arises from germinal center (GC) B cells. Most cases carry the t(14;18) translocation, which drives constitutive BCL2 expression and blocks apoptosis (1). Median survival exceeds ten years, but the course varies widely. The worst outcomes come from two events: histologic transformation to diffuse large B-cell lymphoma (DLBCL), at 2-3% per year (2), and progression within 24 months of first immunochemotherapy (POD24), which affects about 20% of treated patients. FLIPI, the standard prognostic index, uses clinical variables only and says nothing about the biology behind these divergent courses (3).

The microenvironment of FL has long been recognized as a determinant of clinical behavior. Two decades ago, Dave et al. showed that gene-expression signatures from the non-malignant immune cells infiltrating the tumor, rather than from the malignant B cells themselves, correlated with survival in follicular lymphoma. A T-cell-enriched signature (IR-1) marked better outcomes, and a macrophage-and dendritic-cell-enriched signature (IR-2) marked worse (4). Later work tied specific populations to outcome. The architecture of FOXP3+ regulatory T cells predicts survival and transformation (5). CD8+LAG3+ exhausted T cells accumulate in pre-transformation biopsies (6). PD-1+TCF7- cytotoxic T cells track poor survival across nodal B-cell lymphomas (7). Most of this came from dissociated single-cell assays that discard tissue context, or from cohorts too small to test outcome associations. Radtke et al. applied 40-plex imaging to FL lymph nodes and linked enhanced stromal remodeling and distinct stromal communities to high-risk disease and early relapse (8), but examined only a small number of whole sections.

Yet the spatial logic of immune evasion in FL has not been resolved. Normal germinal centers are polarized into light and dark zones, but FL follicles lack this polarity (8,9), and whether they retain any internal organization of their own has been debated. The follicle boundary, where malignant B cells meet the surrounding immune infiltrate, is a distinct transition zone that has drawn little attention. No study has mapped, at single-cell resolution and across a large cohort, where CD8 T cells become exhausted or how regulatory and effector cells distribute across the compartments.

Multiplexed tissue imaging now permits direct examination of these questions. Imaging mass cytometry (IMC) measures dozens of proteins per cell at subcellular resolution without tissue dissociation, allowing cell types, their spatial neighbors, and their functional states to be characterized in situ (10). Two groups have applied such methods to FL. Liu et al. used IMC on 13 paired biopsies obtained at diagnosis and at disease progression and identified peri-follicular immune barriers in early progressors (11). Pelcovits et al. applied spatial transcriptomics and defined four recurrent FL tissue domains (12). Both cohorts were small, which limited their statistical power to associate spatial features with outcome.

Here we report a spatial proteomic atlas of the FL microenvironment. We applied two complementary 39-marker antibody panels to serial sections of four tissue microarrays, generating 4.2 million spatially resolved cells from 156 tumor cores. Of these, 139 cores from 131 clinically annotated patients supported the outcome analyses, and 17 cores from a commercial microarray were used for spatial analyses only. The atlas resolves the neoplastic follicle and its surrounding compartments at single-cell resolution and relates their spatial organization to clinical outcome. It reveals that FL segregates immune evasion into two spatially distinct programs and identifies per-patient CD14 intensity as a prognostic spatial biomarker.

## RESULTS

### Spatial tissue compartments reveal follicular architecture

To characterize the spatial organization of the FL microenvironment, we applied two complementary 39-marker antibody panels to serial sections of four tissue microarrays (**Table S1**). The T-panel targeted T cell subsets and exhaustion markers, and the S-panel targeted myeloid, stromal, and checkpoint molecules. Most antibodies on both panels gave signal adequate for cell-type classification and functional analysis (**Figure S1a,b**). Cellpose hybrid segmentation produced well-separated single cells with a median area of 68 µm² (**Figure S1c,d**; see Methods). The two panels together yielded 4.2 million spatially resolved cells across 156 tumor cores. Of these, 139 cores from 131 patients came from three microarrays constructed at BC Cancer (Vancouver, Canada; A1, B1, C1), and 17 came from a commercial microarray (Biomax) (**Table 1**, **Figure 1a**). All BC Cancer cores were from diagnostic FL biopsies of grade 1 to 3A (**Table 1**), and a subset of patients contributed biopsies from two time points. The Biomax cores lacked clinical annotation and entered the spatial analyses only, not the survival analyses. Hierarchical marker gating identified 15 cell types on the T-panel and 18 on the S-panel, each validated by marker-expression heatmaps (**Figure 1b-g**; see Methods). Macrophages and B cells were larger than T cells, as expected (**Figure S1e**). The major cell-type proportions (40-48% B cells, 17-23% T cells, with CD4 exceeding CD8) agreed with CyTOF and scRNA-seq studies of FL (2,6,8) and differed from normal tonsil (**Figure S1f**). Cross-platform comparison is nonetheless limited by differences in tissue dissociation, antibody panels, and gating.

**Figure 1.**
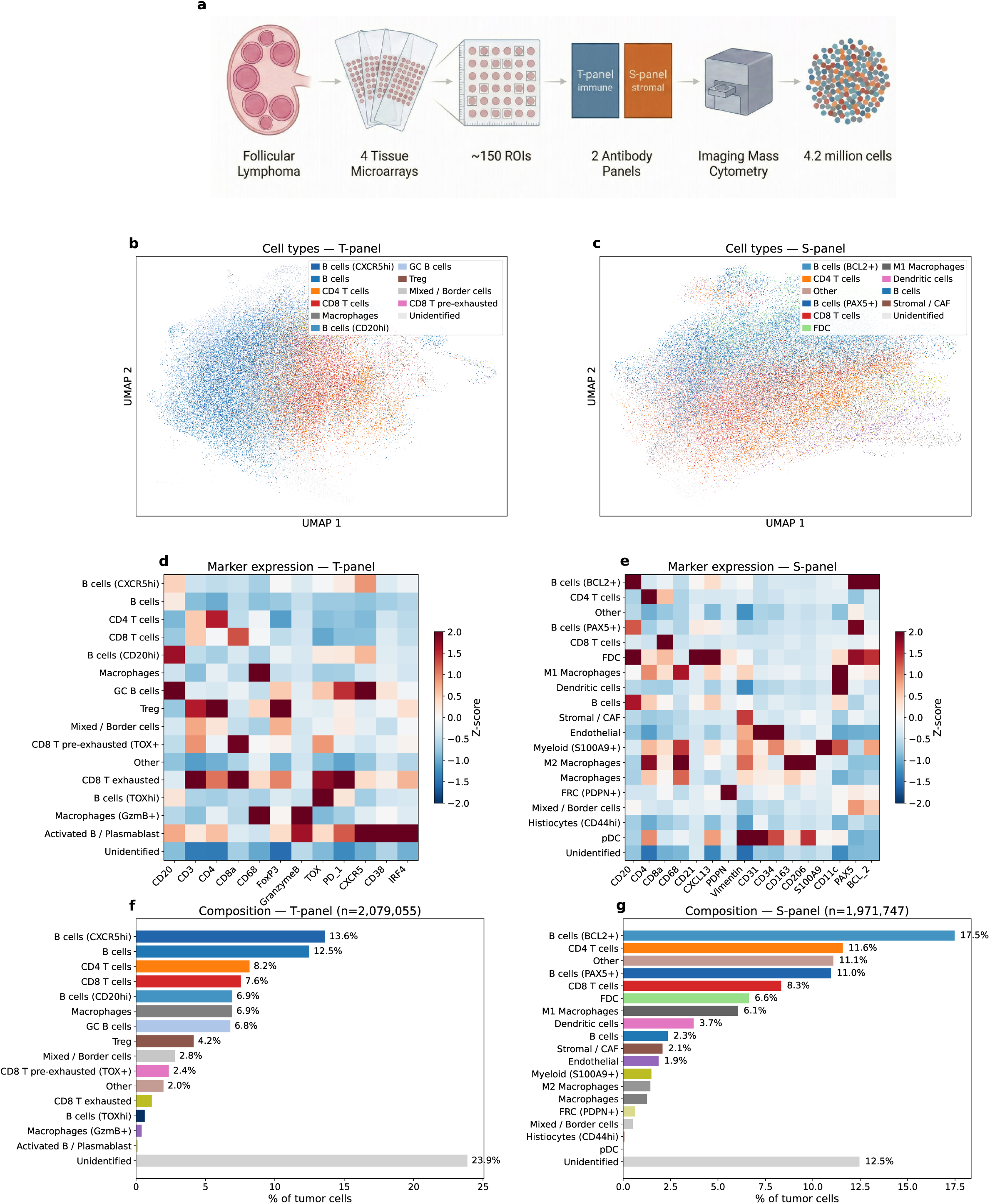
Dataset overview and cell type annotation. (a) Study design: follicular lymphoma tissue on 4 TMAs, ∼150 tumor ROIs, imaged by mass cytometry with T-panel (immune) and S-panel (stromal/myeloid), yielding ∼4.2M cells. (b-c) UMAP embeddings colored by cell type for T-panel and S-panel; includes all cells (FL tumor cores plus control tissues on the same TMAs: tonsil, prostate, kidney, spleen, adrenal). (d-g) Marker expression heatmaps and composition profiles restricted to FL tumor cores. (d-e) Mean marker expression heatmaps (z-scored) validate cell type identity. Apparent CD20/PAX5/BCL2 signal on the S-panel FDC row reflects segmentation spillover from surrounding dense B cell neighborhoods, not B-cell identity; FDC calls are defined by the FDC-intrinsic markers CD21 and CXCL13 with an explicit ‘CD20<CD21’ guard (see Methods, *Cell type annotation*, for the quantitative spillover analysis). (f-g) Cell type composition across tumor cores.

**Table 1.** Patient cohort summary. Clinical and demographic characteristics of the 131 patients with full clinical annotation, split by treatment status (Treated, Observation). Columns show counts and percentages for sex, age at diagnosis, Ann Arbor stage, FLIPI score, grade, and transformation events. Median follow-up and event counts (progression/relapse, death, transformation) reported for each group.

|  | All (N=131) | Treated (N=89) | Obs. (N=42) |
| --- | --- | --- | --- |
| ***Demographics*** |  |  |  |
| Male / Female | 68 / 63 | 45 / 44 | 23 / 19 |
| Age, median (IQR) | 58 (50-64) | 58 (49-64) | 58 (53-63) |
| ***Disease*** |  |  |  |
| Grade 1 | 63 (48%) | 41 (46%) | 22 (52%) |
| Grade 2 | 44 (34%) | 29 (33%) | 15 (36%) |
| Grade 3A | 21 (16%) | 16 (18%) | 5 (12%) |
| Stage III-IV | 108 (82%) | 77 (87%) | 31 (74%) |
| FLIPI High | 46 (35%) | 43 (48%) | 3 (7%) |
| Transformation | 25 (19%) | 19 (21%) | 6 (14%) |
| ***Outcomes*** |  |  |  |
| OS events | 52 (40%) | 38 (43%) | 14 (33%) |
| PFS events | 93 (71%) | 60 (67%) | 33 (79%) |
| Median OS (yr) | 13.7 | 12.3 | 14.5 |
| Median PFS (yr) | 8.6 | 7.5 | 9.9 |

Several quality-control analyses supported these annotations. On serial sections the two panels aligned closely: DNA-based registration confirmed tissue overlap (**Figure S2a**), and the T- and S-panel measurements were spatially concordant, as exemplified by their smoothed CD20-expression maps in a representative core (r=0.90; **Figure S2b**). Cell-type fractions in each ROI agreed across panels for CD4 T, CD8 T, and myeloid cells (Pearson r=0.63-0.75; **Figure S2c,d**). B cell fractions agreed less well overall (r=0.46) but reached r=0.75 in high-quality ROIs (**Figure S2e**). Finally, targeted sequencing of 15 recurrently mutated genes showed no association between mutation status and cell-type composition after correction for multiple testing (all q>0.07; **Figure S3**). This indicates that the contribution of tumor genetics to the microenvironment composition is limited.

We postulated that the FL microenvironment is organized into discrete spatial compartments with distinct cellular composition and function. To test this, we used unsupervised tissue architecture graph (UTAG) analysis (13) to group cells into spatial communities defined by the cell types in their local neighborhood (see Methods), yielding nine recurrent, biologically interpretable tissue compartments (**Figure 2a,b**). Cell type composition shifted smoothly from B cell-dominated follicular compartments (93.8% B cells at the GC core) to T cell-dominated interfollicular zones (**Figure 2c**). Compartment examples across representative ROIs illustrate the spatial diversity of these domains (**Figure S4a-i**). The S-panel resolved additional stromal compartments, including the FDC network zone and stromal/CAF zone (**Figure S5a-i**), and the FDC network zone tracked faithfully with raw CD21 IMC signal (**Figure S6a-d**). Together, these analyses establish a spatially resolved compartment framework (nine on the T-panel, six on the S-panel) that provides the foundation for investigating immune cell function across the FL microenvironment.

**Figure 2.**
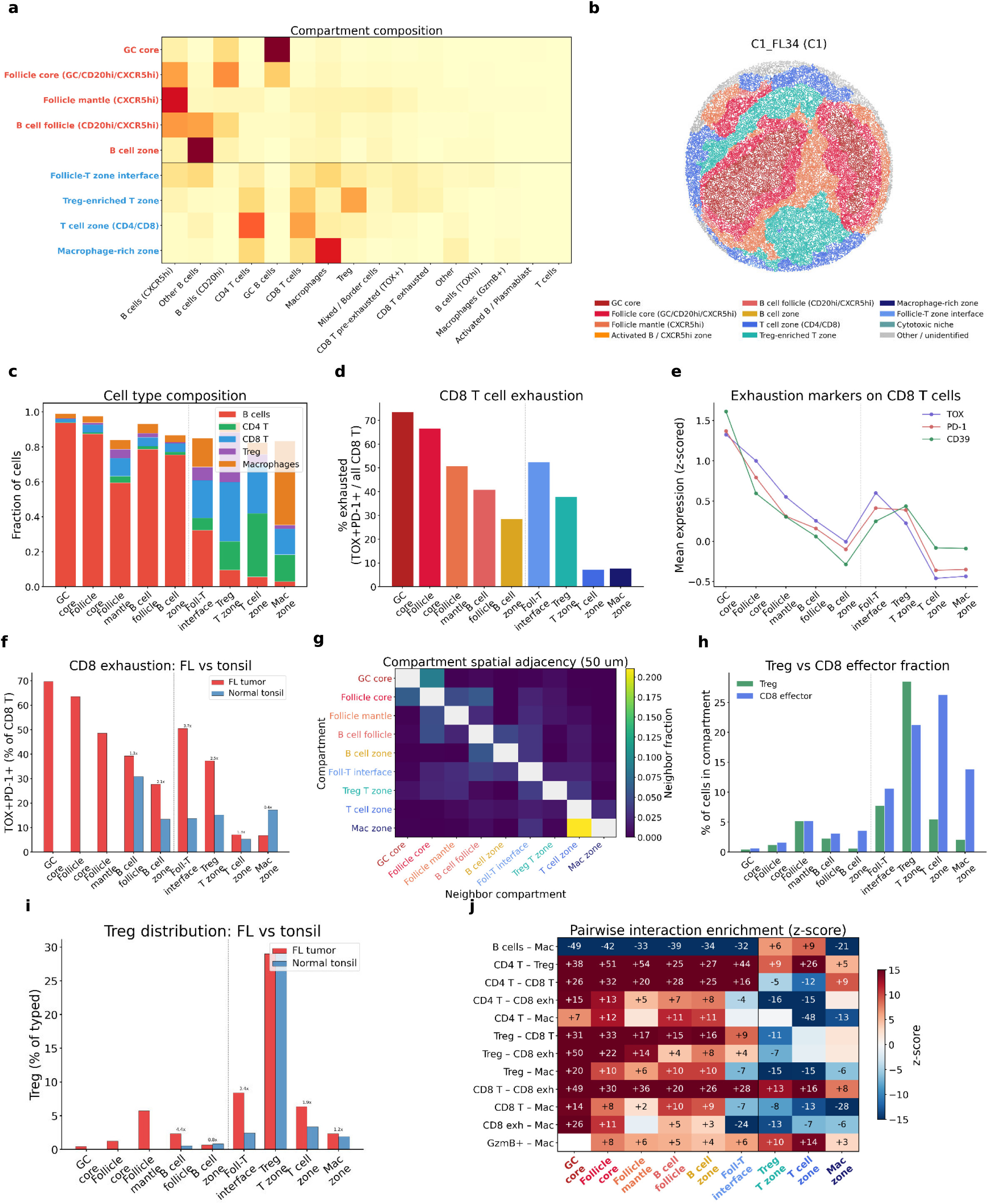
Spatial tissue compartments and follicular biology. UTAG tissue domain analysis using one-hot cell-type features (max_dist=50 µm). (a) T-panel cell type composition per compartment (heatmap; red sidebar = follicular, blue = interfollicular). (b) Representative ROI (C1_FL34) colored by compartment. (c) Cell type composition gradient across 9 compartments (follicle → T zone). (d) CD8 T cell exhaustion fraction (TOX+PD-1+): 73.6% at GC core vs 7.2% in T zone. (e) Exhaustion marker expression (TOX, PD-1, CD39) on CD8 T cells. (f) FL vs tonsil CD8 exhaustion comparison. (g) Compartment spatial adjacency (row-normalized fraction of each compartment’s neighbors within 50 µm, diagonal masked): off-diagonal mass concentrates on gradient-adjacent compartments, ordering them concentrically. (h) Treg vs CD8 effector fraction: Treg dominates follicular zones. (i) FL vs tonsil Treg distribution: FL-specific Treg enrichment. (j) Pairwise interaction enrichment (permutation z-scores, K=10, 200 perms): Mac–CD8 T and Mac–exhausted-CD8 T enriched follicularly (GC core z=+15, +27), depleted in T cell zone (z=−13, −7).

### Immune exclusion and exhaustion follow a compartment-dependent gradient

Having established distinct tissue compartments, we next asked whether immune cell distribution and functional state vary across the follicular-to-interfollicular axis. CD8 T cells constituted only 2.0% of cells at the GC core compared to 28.3% in interfollicular zones (**Figure 2c**), confirming immune exclusion from neoplastic follicles. Among the CD8 T cells that did infiltrate follicular zones, exhaustion was pervasive: co-expression of TOX and PD-1 decreased sharply from 73.6% at the GC core to 7.2% in the T cell zone (**Figure 2d**), with exhaustion markers TOX, PD-1, and CD39 all following the same gradient (**Figure 2e**). This exhaustion pattern was specific to FL: comparison with normal tonsil controls revealed similar anatomical exclusion of CD8 T cells from follicles, but exhaustion rates were substantially lower in tonsils across all compartments (**Figure 2f**). These compartments are spatially ordered rather than interspersed: at 50 µm resolution each compartment borders predominantly its neighbors along the follicular-to-interfollicular axis, so the mean position of a compartment’s spatial neighbors increases monotonically with its own position (**Figure 2g**). Treg and CD8 effector cells showed distinct spatial distributions: Tregs peaked at the follicle-T zone interface and in the Treg-enriched T zone, while CD8 effectors concentrated in the T cell zone and macrophage-rich zone (**Figure 2h**). The FL-specific Treg enrichment was confirmed by comparison with tonsil controls (**Figure 2i**). Among FoxP3+ cells, the fraction with high CXCR5 expression consistent with a T follicular regulatory (TFR) phenotype (14) followed a steep spatial gradient, from 70% at the GC core to 17% at the follicle-T zone interface and under 2% in the T cell zone (**Figure S7a**). FoxP3+ cells within follicles are therefore predominantly TFR, whereas those forming the boundary barrier and the T zone are predominantly classical Tregs. Permutation-based pairwise interaction analysis (K=10 neighbors, 200 permutations) revealed compartment-specific cell-cell interaction patterns: macrophage–CD8 T and macrophage–exhausted-CD8 T interactions were enriched at the GC core (z=+15 and +27 respectively) and depleted in the T cell zone (z=−13 and −7) and macrophage-rich zone (z=−28). Macrophage–CD4 T and macrophage–Treg pairs followed the same follicular-enrichment pattern (GC core z=+7 and +20), consistent with rare follicular macrophages preferentially neighboring infiltrating T cells. B cell–macrophage pairs were depleted in B-rich follicular compartments (GC core z=−49, follicle core z=−42), reflecting the low absolute frequency of macrophages among the B-cell background rather than active avoidance (**Figure 2j**). Because Tfh cells are essential for maintaining the germinal center reaction and have been implicated in FL pathogenesis (9), we examined their spatial distribution. PD-1-high (>1.5) T follicular helper (Tfh) cells were densest at the follicle mantle and the Treg-enriched T zone, rather than at the GC core (**Figure S7b**). The small number of Tfh that did reach the GC core were predominantly in the PD-1-high state, indicating that the innermost follicular environment selects for the most activated phenotype. Both the peripheral and GC-core Tfh expressed elevated CXCR5, PD-1, and TOX compared to non-Tfh CD4 T cells (**Figure S7c**). The peripheral Tfh remained predominantly CXCR5+, confirming that they are bona fide CXCR5+ Tfh rather than a CXCR5-negative peripheral population. Permutation-based neighborhood enrichment (K=10, 500 permutations) at the GC core showed that PD-1-high Tfh were consistently in close proximity to malignant B cells (z=+10.6), CD4 T cells (z=+11.3), and FoxP3+ cells (z=+22.3; **Figure S7d**). Because GC-core FoxP3+ cells are predominantly TFR (**Figure S7a**), these neighbors are TFR rather than classical Tregs. Spatial mapping of representative cores showed that PD-1-high Tfh form localized clusters at the follicle rim and follicle-T zone interface rather than distributing uniformly throughout the follicle interior (**Figure S7e-g**). Collectively, these findings reveal that the FL follicle operates as an immune-privileged zone where CD8 T cells are both excluded and functionally impaired, separated from the effector-rich interfollicular compartment by a Treg-enriched boundary.

### Histologic grade is accompanied by macrophage decompartmentalization and quantifiable follicle boundary irregularity

Histologic grade is a standard histological descriptor of FL, but its single-cell spatial correlates have not been mapped. We asked how the microenvironment changes across grade, and whether the spatial features that predict outcome (next section) also track it. As an internal consistency check, the Ki-67⁺ B-cell fraction rose from 7.5% in FOLL1 to 16.3% in FOLL3A (Kruskal–Wallis p=0.007; **Figure S8a**), confirming that our single-cell typing recovers the centroblast-count gradient that defines grade.

The reorganization with grade went well beyond proliferation. Macrophage populations expanded. M1 macrophages and S100A9⁺ MDSC-like cells roughly tripled between FOLL1 and FOLL3A (p=0.001 and p=0.020; **Figure S8b,c**). On the independent T-panel, which does not resolve myeloid subsets, total macrophages doubled across the same range (p=0.004; **Figure S8d**), confirming the expansion on a second panel. These extra macrophages did not crowd into the existing micro-niches, instead redistributed across the tissue. At the scale of single-cell neighborhoods, the tendency of macrophages to cluster next to one another declined (Ripley’s L, a spatial measure of same-type clustering, fell 32% at 25 µm from FOLL1 to FOLL3A; p=0.005; **Figure S8f**). At the tissue scale, the more abundant macrophages formed a discrete macrophage-rich UTAG compartment that was effectively absent at lower grade (p=0.002; **Figure S8e**). The follicle outline also grew less regular: mean compactness (the isoperimetric ratio 4πA/P², equal to 1 for a perfect circle) declined from 0.46 to 0.31 across grades (p=0.021; **Figure S8g**). Two representative cores illustrate the cohort-wide pattern (**Figure S8h-i**): a low-grade core with several discrete, round follicles and sparse macrophages, and a grade-3A core with a single, irregular follicular mass infiltrated by abundant, dispersed macrophages.

### CD14 is the strongest predictor of clinical outcomes

To determine whether the compartmentalized organization of the FL microenvironment carries clinical significance, we screened cell-type fractions against progression-free survival (PFS) and overall survival (OS) in treated patients (n=116; observation-only patients excluded, see Methods). M2 macrophages and FDCs were the strongest cellular predictors of both endpoints (**Figure 3a,b**). S100A9+ MDSC-like myeloid cells ranked among the strongest predictors of histologic transformation (**Figure 3c**). Among individual markers, the per-ROI mean CD14 staining intensity on the S-panel was the strongest predictor of PFS (HR=1.59, P<0.0001; **Figure 3d**). This metric averages per-cell CD14 expression across the ROI rather than counting CD14+ cells, and CD14 is expressed by both macrophages and FDCs (characterized in the following section). A median split separated patients into worse and better PFS groups (**Figure 3e**). CD14 withstood progressive covariate adjustment, remaining significant after adding FLIPI, then grade, then stage and age (**Figure 3f**). In the full clinical model it was the only independent predictor of PFS (HR=1.50, P=0.0007). FLIPI, grade, stage, and age were not significantly associated with PFS (**Figure 3g**). Combining CD14 intensity with FLIPI predicted early progression (POD24) with an in-sample area under the curve of 0.77 (**Figure 3h**), above FLIPI alone (0.68) or CD14 alone (0.64). CD14 intensity therefore captures prognostic microenvironment biology that conventional clinical scoring does not.

**Figure 3.**
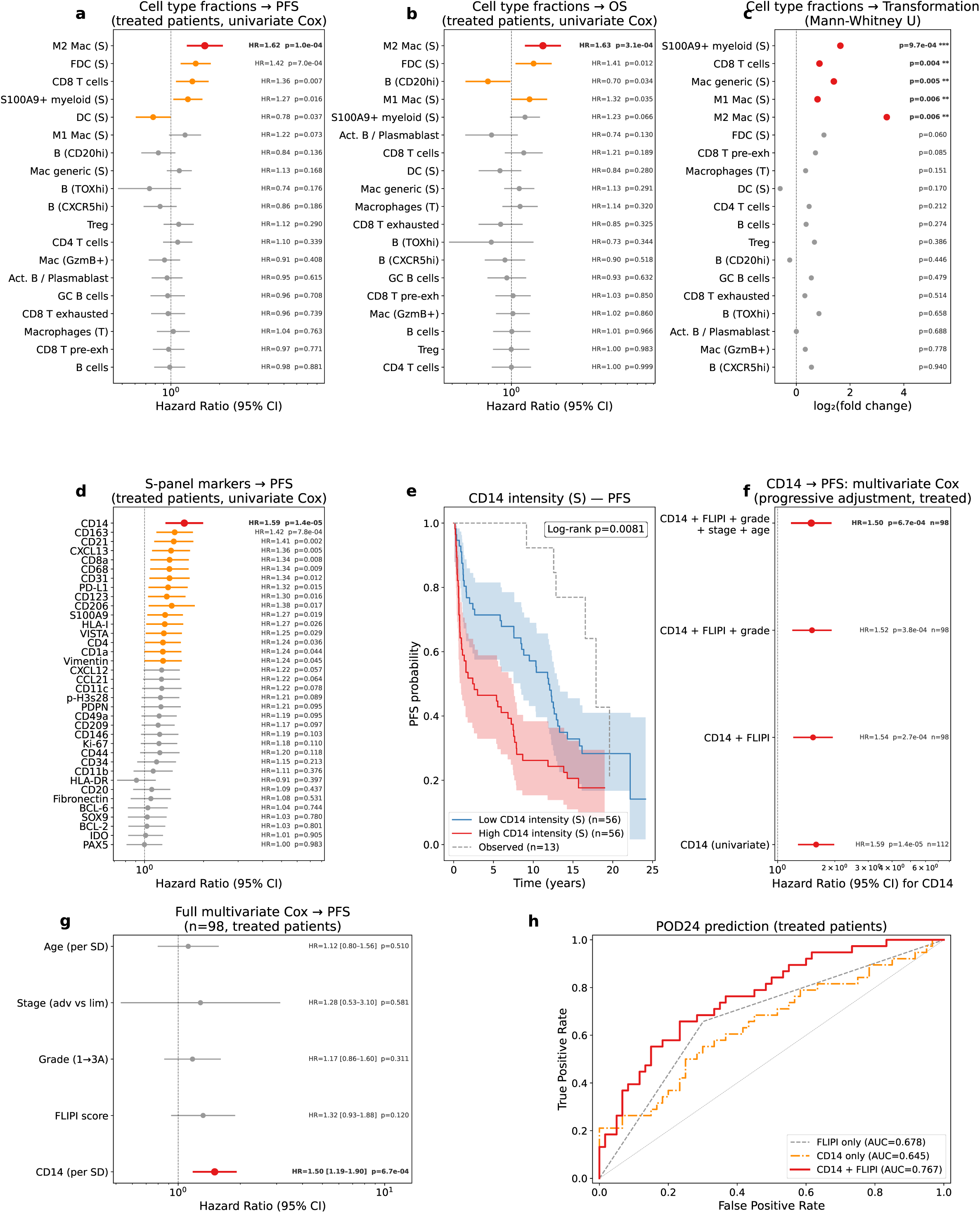
Survival analysis, biomarker discovery, and multivariate validation. Comprehensive survival screen and CD14 biomarker validation in treated patients. (a) Cell-type fraction forest — PFS (univariate Cox, HR per SD): M2 Mac and FDC predict shorter PFS. (b) Cell-type fraction forest — OS: M2 Mac, FDC, S100A9+ predict shorter OS. (c) Transformation forest (Mann-Whitney, all patients): S100A9+ (3.4x), M1 Mac (1.9x), M2 Mac (15.6x) enriched in transformers. (d) S-panel marker forest — PFS: CD14 (HR=1.59, P<0.0001) is the strongest predictor. (e) KM: CD14 high vs low splits PFS. (f) Multivariate Cox: CD14 HR remains significant across progressive adjustment (univariate → +FLIPI → +grade → +stage+age; full model HR=1.50, P=0.0007). (g) Full model forest: CD14 is the only independent predictor of PFS (FLIPI P=0.088, grade/stage/age all NS). (h) POD24 prediction: CD14+FLIPI (AUC=0.77) vs FLIPI-only ROC.

### CD14-high FDCs are activated stromal organizers of the intrafollicular niche

To identify the cellular origin of the CD14 prognostic signal, we examined CD14 expression across cell types. CD14 expression was highest on myeloid cells but unexpectedly was also elevated on FDCs (15), the second-highest expressing cell type (**Figure 4a**). We defined CD14-high FDCs as those above the 75th percentile of CD14 intensity among all FDCs (referred to as CD14+ when discussing scRNA-seq binary detection). Because CD14+ cells in IMC could reflect proximity-dependent signal spillover (signal bleed from adjacent cells) from neighboring myeloid cells, we decomposed the CD14+ population: myeloid cells accounted for 30%, FDCs for 25%, with the remainder distributed across other cell types (**Figure S9a**). To ascertain whether the CD14 on FDCs was merely spillover, we asked whether a cell’s CD14 positivity depended on its distance from the nearest myeloid cell. This dependence was weak (Spearman rho=-0.115), indicating that spillover contributes to but does not fully explain the CD14 signal on FDCs. Independent validation using a published FL scRNA-seq atlas (Han et al., 2022 (16); an independent cohort unrelated to our IMC samples) confirmed that FDCs express CD14 mRNA, with 19% of FDCs scoring positive (**Figure S9b**), and FL FDCs expressed significantly more CD14 than normal tonsil FDCs (P=1.4e-44; **Figure S9c**).

**Figure 4.**
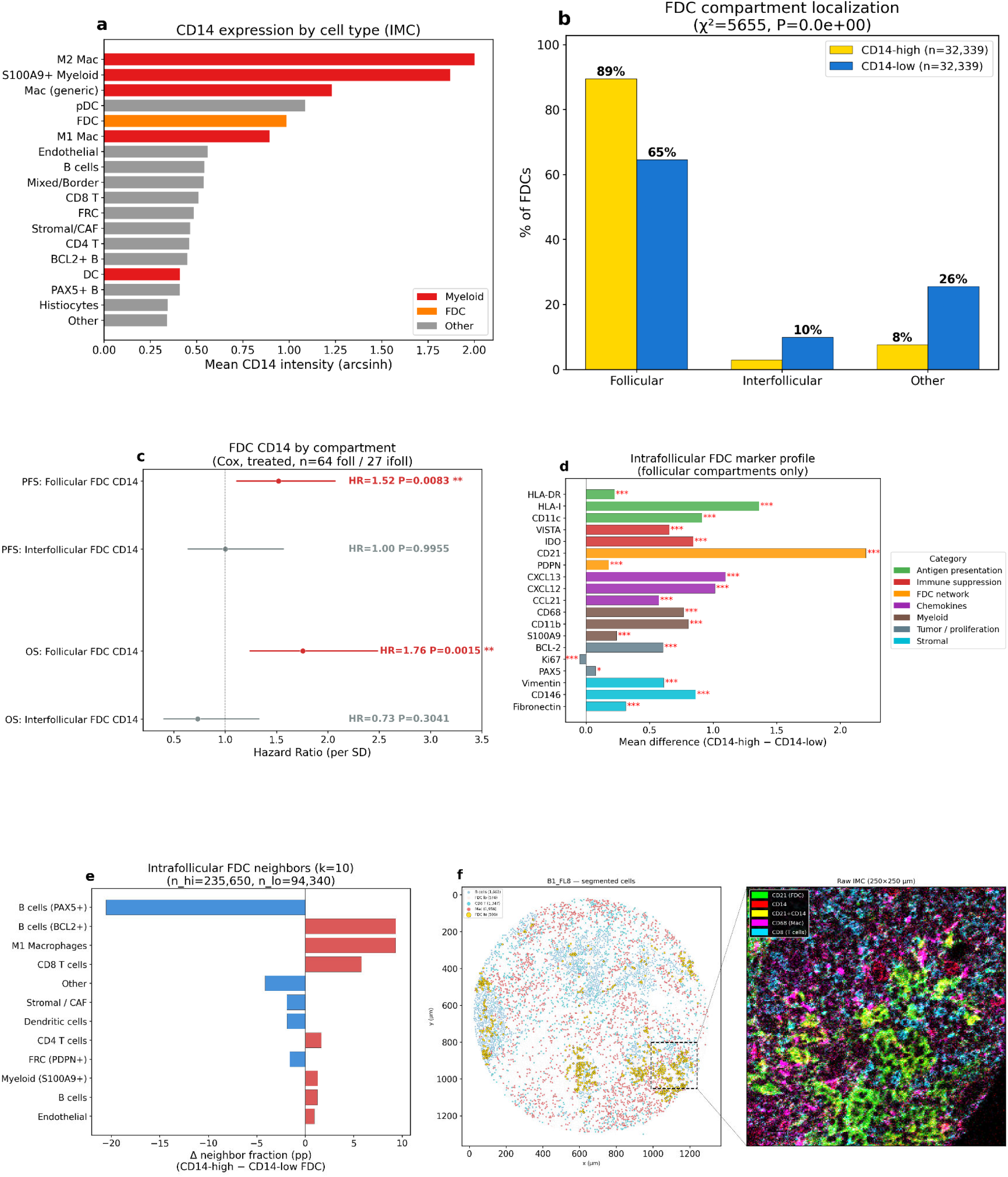
CD14+ FDCs: discovery, characterization, and tumor support. (a) CD14 expression by cell type (IMC): FDC is second highest after myeloid. (b) Compartment localization: CD14-high FDCs 89% follicular vs 65% for CD14-low (chi-sq=5655, P<0.001). (c) Compartment-specific survival: follicular FDC CD14 predicts PFS (HR=1.52, P=0.008) and OS (HR=1.76, P=0.0015); interfollicular FDC CD14 does not (PFS HR=1.00, OS HR=0.74, both NS). (d) Intrafollicular marker profile (follicular compartments only): CD14-high FDCs upregulate CD21 +2.2, HLA-I +1.4, CXCL13 +1.1, and VISTA. (e) Intrafollicular neighbors (k=10): CD14-high FDCs enriched for BCL2+ B (+9.3pp), M1 Mac (+9.3pp), CD8 T (+5.8pp); depleted for PAX5+ B (−20.6pp). (f) Representative tissue microenvironment (B1_FL8): segmented cell scatter with raw IMC composite inset (CD21=green, CD14=red, CD68=magenta, CD8=cyan; CD21+CD14 overlap appears yellow) showing CD14-high FDCs co-localizing with macrophages and CD8 T cells.

CD14-high FDCs were concentrated in follicular compartments (89% follicular versus 65% for CD14-low FDCs, chi-squared P<0.001; **Figure 4b**). Only intrafollicular FDC CD14 predicted outcome: it was associated with shorter PFS (HR=1.52, P=0.008) and OS (HR=1.76, P=0.0015), whereas interfollicular FDC CD14 showed no association (**Figure 4c**). Within follicular compartments, CD14-high FDCs expressed more CD21, HLA class I, CXCL13, and VISTA than their CD14-low counterparts (**Figure 4d**). Re-analysis of the Han et al. scRNA-seq atlas (16) showed that CD14+ FDCs carried 1.6-fold more total mRNA per cell than CD14- FDCs (median 3,009 versus 1,922 UMIs, P=6.0e-7; **Figure S9d**), consistent with a transcriptionally active state. These CD14+ FDCs clustered with other FDCs rather than with myeloid cells and did not express myeloid lineage genes. CD14 therefore marks an activated FDC state, not myeloid contamination.

CD14-high FDCs formed spatially contiguous communities rather than scattering individually through the follicle: 60% of their ten nearest neighbors were themselves CD14-high FDCs (vs 2% for CD14-low FDCs, Δ=+57pp; data not shown), and CD14-low FDCs likewise clustered with other CD14-low FDCs (30% vs 1%, Δ=+30pp; data not shown), indicating that the two FDC states occupy distinct local niches. Among non-FDC neighbors, CD14-high FDCs were preferentially surrounded by BCL2+ B cells, M1 macrophages, and CD8 T cells compared to CD14-low FDC neighborhoods (**Figure 4e**). Follicular B cells near CD14-high FDCs showed higher Ki-67 expression than B cells near CD14-low FDCs (mean 0.483 vs 0.301, P=2.9e-32; **Figure S9e**) and higher MHC class I and class II expression (**Figure S9f**), suggesting that these activated FDCs support tumor proliferation and modulate antigen presentation-associated signaling. scRNA-seq showed that CD14+ FDCs upregulate the B cell survival signals TGF-beta1 and IL-6, with a modest trend toward elevated APRIL (**Figure S9g**). Raw IMC composite images confirmed this niche directly (**Figure 4f**). Within a representative follicle (B1_FL8), CD21+ FDCs (green) and CD14 (red) co-localized at the pixel level, appearing yellow where they overlapped. CD68+ macrophages (magenta) and scattered CD8 T cells (cyan) occupied the same field. This matched the segmented-cell map, in which CD14-high FDCs sat embedded within clusters of B cells, macrophages, and CD8 T cells. At the tissue level, FDCs were the dominant source of CXCL13 and CCL21, whereas BAFF and APRIL were predominantly myeloid-derived (**Figure S9h**), consistent with recent IMC findings in FL from our group (17). CXCL13 expression correlated with CD21 meshwork density across ROIs (rho=0.56; **Figure S9i**). An integrated signaling heatmap confirmed the compartmentalized checkpoint landscape, with VISTA peaking on M2 macrophages and CXCL13 on FDCs (**Figure S9j**). These data establish CD14-high FDCs as disease-specific activated stromal cells that organize the intrafollicular niche, support tumor B cell survival, and as noted earlier independently predict clinical outcomes.

### The myeloid ecosystem is compartmentalized and dominated by VISTA

Given the prominence of myeloid populations in the survival and transformation analyses, we characterized the spatial organization and functional specialization of myeloid subtypes in detail. The S-panel resolved three major myeloid subtypes — M1 macrophages, M2 macrophages, and S100A9+ MDSC-like cells — plus dendritic cells and pDCs analyzed separately. The three subtypes had distinct functional marker profiles (**Figure 5a**): M2 macrophages expressed the highest levels of VISTA and CCL21; S100A9+ cells were defined by S100A9, CD14, and CD11b; and M1 macrophages showed intermediate expression of most markers.

**Figure 5.**
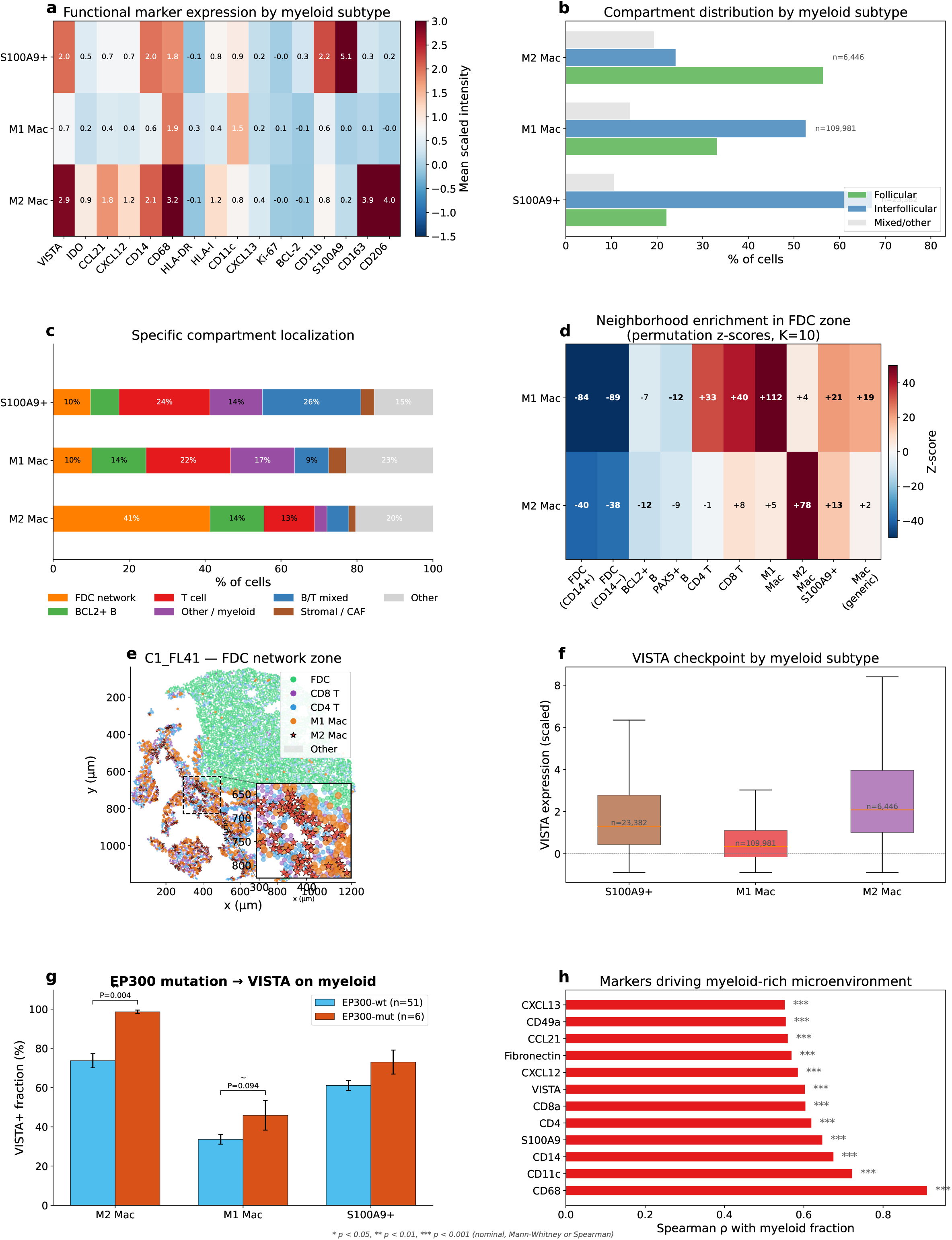
Myeloid ecosystem: compartmentalization and spatial interactions. Spatial organization and functional specialization of M1, M2, and S100A9+ myeloid subtypes. (a) Functional marker profiles. (b) Compartment distribution: M2 Mac uniquely follicular (56.5%); M1, S100A9+ predominantly interfollicular. (c) Specific compartment localization: M2 Mac concentrates in the FDC network zone (41%); M1 and S100A9+ distribute across T cell and other interfollicular zones. (d) Permutation-based neighborhood z-scores (k=10 nearest, 200 perms) within the FDC network zone; z-scores clipped at ±50 for visibility (M1/M2 self-enrichment reaches +83/+122). (e) Myeloid-lymphocyte distances. (f) VISTA checkpoint expression by myeloid subtype. (g) EP300 mutation associated with VISTA upregulation, strongest in M2 Mac (P=0.004). (h) Marker drivers of myeloid-rich microenvironment. See Figs S10-S12 for detailed M2 Mac, S100A9+, and VISTA analyses.

These subtypes partitioned spatially in the tissue. M2 macrophages were the only myeloid subtype enriched in follicular zones (56.5% follicular; **Figure 5b**), and at finer resolution concentrated specifically in the FDC network zone (41% of all M2 Mac; **Figure 5c**). Within the FDC network zone, permutation-based neighborhood analysis (K=10, 200 perms) showed that M2 macrophages formed strongly self-clustered pockets (z=+83) enriched for S100A9+ cells (z=+16) and CD8 T cells (z=+8), but were depleted as direct FDC neighbors (z=-40 for CD14+ FDC, z=-34 for CD14-) and from dense B cell clusters (**Figure 5d**). A representative ROI (**Figure 5e**, **Figure S10a**) illustrates this pattern at the compartment scale: M2 macrophages and CD14+ FDCs both populate the FDC network zone but remain spatially separated at the single-cell level, and a subset of M1 macrophages was also present in this zone (weakly co-localized with M2 Mac, z=+5), consistent with their enrichment as non-FDC neighbors of CD14-high FDCs in **Figure 4e**. At the population scale, however, M1 macrophages distributed predominantly interfollicularly (55% interfollicular vs 30% follicular), and S100A9+ cells were even more strongly interfollicular (67%) (**Figure 5b**), occupying T cell zones and mixed compartments (**Figure 5c**).

Functionally, VISTA was the dominant checkpoint across all myeloid subtypes, expressed at highest levels on S100A9+ cells and M2 macrophages, while PD-L1 signal was near-absent (**Figure 5f**), consistent with the low PD-L1 dynamic range observed on our panels across all TMAs (**Figure S1a,b**). Patients carrying EP300 mutations, a recurrent alteration in FL, showed upregulated VISTA on myeloid cells — most strongly on M2 macrophages (P=0.004; **Figure 5g**) — linking tumor genetics to microenvironment remodeling. Per-ROI marker correlation identified CD68, CD11c, S100A9, and VISTA as the strongest drivers of myeloid-rich microenvironments; CXCL13 and CD49a specifically marked the FDC niche (**Figure 5h**).

Because S100A9+ cells were among the strongest predictors of transformation (**Figure 3c**), we examined their biology in more detail. S100A9+ cells in interfollicular zones were surrounded by M1 macrophages, CD8 T cells, and B cells (**Figure S11a,b**), and per-ROI S100A9+ fraction correlated positively with M1 macrophages (rho=+0.53) and cytotoxic CD8 T cells (**Figure S11c**) — marking inflamed microenvironments. The rare S100A9+ cells that reached the follicle expressed higher VISTA than their interfollicular counterparts (**Figure S11d**), consistent with an elevated suppressive phenotype within the tumor niche. In the Han et al. FL scRNA-seq atlas (16), S100A9+ cells expressed the calprotectin program (S100A8, S100A9, S100A12) and showed concordance with the IMC protein markers CD14, CD11b, and VISTA (**Figure S11e**).

Integrating these data, the three myeloid populations are spatially and functionally complementary: M2 macrophages are follicle-associated, co-localize with CD14-high FDCs within the FDC network zone while remaining spatially separated from individual FDCs at the single-cell level, and express the highest levels of VISTA — they are part of the intrafollicular immune-privileged niche. M1 macrophages are interfollicular, more inflammatory in phenotype, and occupy the interfollicular zone alongside S100A9+ cells and CD8 T cells. S100A9+ MDSC-like cells are overwhelmingly interfollicular and are the strongest myeloid predictor of transformation; the small follicular subset carries disproportionately high VISTA. Together, VISTA on M2 macrophages and S100A9+ cells — rather than PD-L1 — defines the dominant immune checkpoint axis in the FL myeloid compartment, with EP300 mutations further amplifying VISTA expression on M2 macrophages.

The Han et al. scRNA-seq atlas (16) characterized the VISTA checkpoint landscape at the transcript level. VISTA+ myeloid cells co-expressed the M2-macrophage markers CD163 and TGF-beta1 along with TIM-3 and Galectin-9, indicating an M2-skewed, immunosuppressive phenotype (**Figure S12a**), and 94% carried two or more additional checkpoints (**Figure S12b**). Across myeloid and FDC populations, VISTA, TIM-3, and Galectin-9 were the most broadly expressed cell-surface checkpoint molecules (**Figure S12c**). At the transcriptome level, VISTA mRNA predominated over PD-L1 in myeloid cells (56% versus 12% positive), confirming the IMC finding that VISTA, not PD-L1, is the operative myeloid checkpoint in FL (**Figure S12d**).

IMC protein data reinforced this hierarchy. Per-cell VISTA+ fractions were highest on follicular M2 macrophages (91%) and S100A9+ cells (88%), and lower but still elevated on CD14+ FDCs (32%, versus 15% for CD14- FDCs) (**Figure S12e**). Because CD14+ FDCs are abundant, they constituted the largest VISTA+ population in the follicular compartment (**Figure S12f**), placing them in frequent VISTA-mediated contact with infiltrating T cells. This VISTA load was specific to FL: per-cell mean VISTA was 3.8-fold higher on M2 macrophages, 3.4-fold higher on S100A9+ cells, and 2.1-fold higher on CD14+ FDCs than in normal tonsil (per-ROI Mann-Whitney p=1.7e-4, 5.5e-4, and 1.4e-2), whereas VISTA on CD14- FDCs did not differ from tonsil (**Figure S12g**). The populations that drive the VISTA-based evasion architecture (M2 macrophages, S100A9+ cells, and CD14+ FDCs) therefore upregulate VISTA in a disease-specific manner, while CD14- FDCs remain at tonsillar baseline. Although the myeloid populations are spatially partitioned between follicular and interfollicular zones, VISTA is expressed across both (on follicular M2 macrophages and CD14-high FDCs, and on interfollicular M1 and S100A9+ cells). It is therefore the checkpoint shared by both immune-evasion programs rather than the feature that separates them.

### Agent-based modeling predicts synergy between anti-VISTA and bispecific CD20-CD3 therapy

To test the therapeutic implications of this architecture, we built an agent-based model from our IMC data, encoding the major cell types, the concentric zones, and the suppressive interactions between them (Methods; **Table S2**). The model reproduced several clinical observations in FL. Anti-VISTA monotherapy lifted myeloid suppression but did not clear the tumor on its own. Rituximab monotherapy produced a deep, durable response (89% reduction, nadir 66 cells at t=777) followed by slow CD20-negative relapse (final ∼155 cells; **Figure 6e-f**), mirroring the 1-2 year progression-free survival seen with rituximab monotherapy. Bispecific CD20-CD3 monotherapy cleared the tumor eventually but slowly (t=766; **Figure 6g-h**). Anti-VISTA added to rituximab remained limited by CD20-negative escape, because rituximab is blocked by CD20 loss and anti-VISTA cannot by itself recruit T cells to the CD20-negative residual. Anti-VISTA combined with bispecific CD20-CD3 cleared the tumor nearly twice as fast as the bispecific alone (t=405 versus t=766; **Figure 6k-l**). In that combination the bispecific bridges T cells to the CD20-positive bulk and retains partial activity against CD20-low cells, while anti-VISTA restores CD8 T cell cytotoxicity. Adding rituximab on top gave only a small further gain (t=177; **Figure 6m-n**), unlikely to justify the added toxicity of a three-drug regimen. The model therefore favors anti-VISTA plus a CD20-CD3 bispecific as a two-drug regimen aimed at the intrafollicular evasion architecture (**Figure 6a-n**).

**Figure 6.**
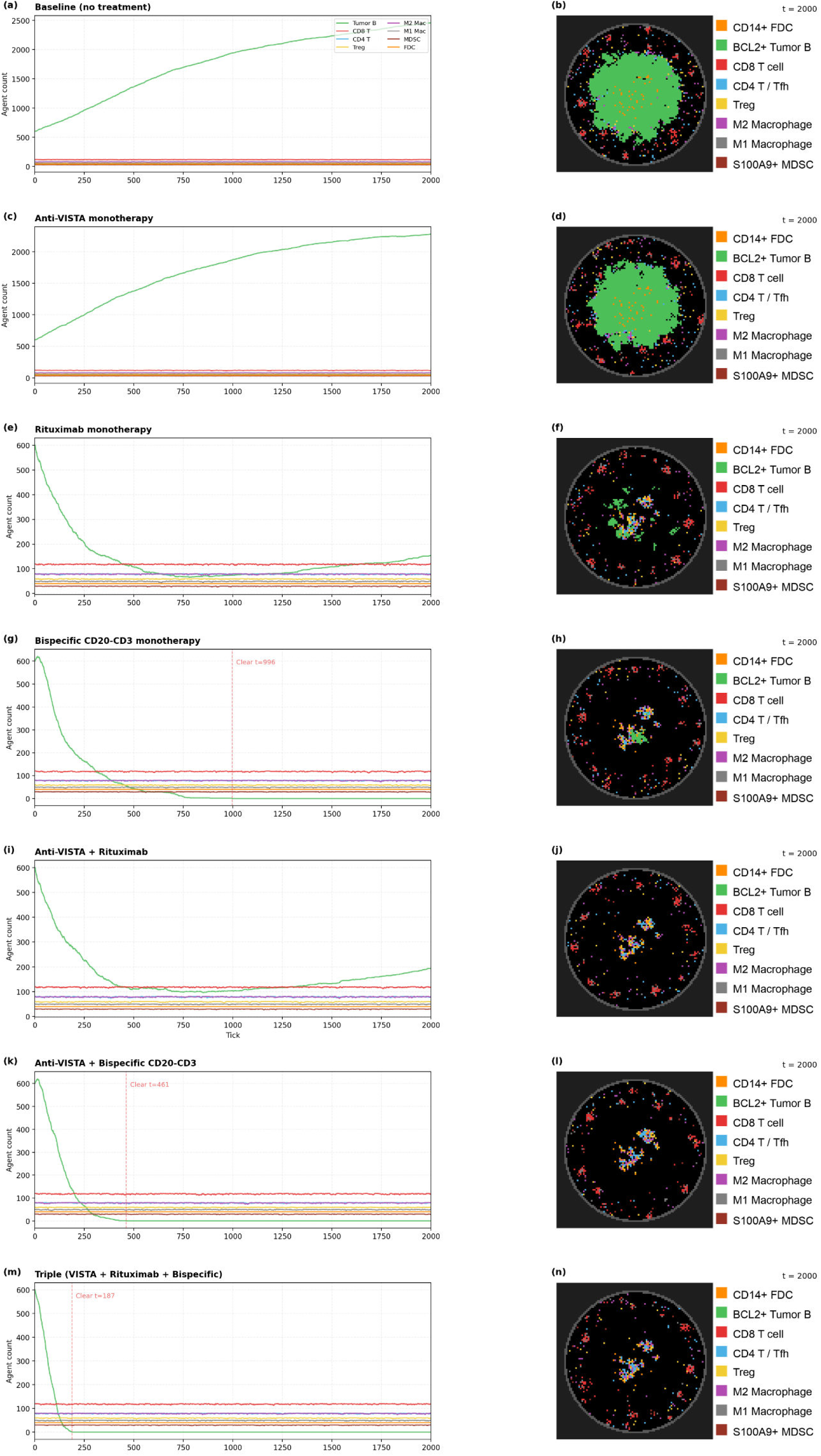
Agent-based model: treatment scenario simulations. Agent-based model (Mesa framework) encoding the dual-compartment evasion architecture: 8 cell types, 4 diffusive chemokine fields, concentric zone geometry (100×100 grid, circular tissue r=45), and a 3% CD20-negative tumor subclone with 20% slower proliferation (fitness cost of CD20 loss). Rituximab is fully blocked by CD20 loss (Weiner 2010, Davis 1999); modern bispecific antibodies retain partial (∼50%) activity against CD20-low cells due to higher per-molecule avidity. Left column: population trajectories over 2000 ticks. Right column: final spatial snapshot (t=2000). (a-b) Baseline: tumor B grows to ∼2460, CD8 T excluded from follicle. (c-d) Anti-VISTA monotherapy: lifts suppression but tumor persists. (e-f) Rituximab monotherapy: deep durable response (nadir ∼66 at t=777), slow CD20-negative relapse (final ∼155). (g-h) Bispecific CD20-CD3 monotherapy: slow clearance (t=766). (i-j) Anti-VISTA + Rituximab: CD20-negative residual (final ∼193), similar to rituximab alone. (k-l) Anti-VISTA + Bispecific CD20-CD3: clearance at t=405, nearly twice as fast as bispecific alone. (m-n) Triple combination: clearance at t=177, modest speed improvement over the two-drug combination. Immune cell turnover (death 0.3%/tick + homeostatic immigration) maintains CD8 at 112-120. Model code: github.com/oelemento/FL-ABM.

## DISCUSSION

Our finding that FL follicles show graded spatial organization — despite lacking the light zone/dark zone polarity of normal germinal centers (8,9) — adds a layer of sub-follicular architecture that appears specific to the neoplastic GC. This extends the growing recognition that germinal center organization is more complex than the classical two-zone model: Kennedy et al. (18) identified additional sub-compartments within the dark zone of normal GCs (a proliferating-cell niche associated with tingible body macrophages), refining the canonical LZ/DZ framework. Our data reveal a distinct, concentric organization that emerges in FL follicles which lack that framework entirely, arising from a reorganized FDC/B cell scaffold rather than the classical centroblast/centrocyte program. The nine compartments we identify define functionally distinct immune microenvironments: a CD8 T cell within the GC core is ten times more likely to be exhausted than one in the T cell zone. This gradient has implications for immunotherapy, as therapies targeting intrafollicular T cells must overcome not only checkpoint signaling but also the physical and regulatory barriers that separate follicular and interfollicular zones.

Our spatial findings complement and extend the multi-omic atlas of Radtke et al. (8). Their study identified DC-SIGN+ FDCs and stromal desmoplasia as features of early relapsers using 40-plex imaging of whole lymph node sections. We confirm the importance of FDC heterogeneity and add three dimensions their study did not address: compartment-dependent T cell exhaustion mapping, the identification of VISTA as the dominant myeloid checkpoint, and a prognostically validated spatial biomarker (CD14) tested across 131 patients. Conversely, their finding of enhanced extracellular matrix remodeling in high-risk patients suggests stromal barriers that may compound the immune evasion architecture we describe.

The identification of CD14-high FDCs as activated stromal organizers of the intrafollicular niche builds on the observation by Smeltzer et al. that the follicular pattern of CD14+ cells, assessed by standard IHC, predicts time to transformation (15). Because CD14 is classically a monocyte/macrophage marker, its expression on FDCs could suggest myeloid misclassification. However, CD14-high FDCs co-express CD21 (a definitive FDC marker not found on myeloid cells) and upregulate it relative to CD14-low FDCs (**Figure 4d**), arguing against lineage contamination. Consistent with this, scRNA-seq analysis shows that CD14+ FDCs do not upregulate myeloid lineage genes but instead show elevated overall transcriptional output, indicating an activated stromal state. The compartment-specific survival association, where only intrafollicular FDC CD14 predicts outcomes, aligns with the principle that it is the spatial pattern, not the quantity, of CD14+ cells that matters clinically. These findings are consistent with the broader understanding that FL reprograms its stromal niche: Mourcin et al. showed that FL B cells induce phenotypic and functional remodeling of FDCs and fibroblastic reticular cells via TNF and TGF-beta signaling (19), and Tfh cells provide CD40L and IL-4 signals that support malignant B cell survival within the follicle (20). CD14-high FDCs may represent the most activated state of this reprogrammed stromal scaffold. This finding also provides a candidate spatial explanation for the IR2 prognostic signature described by Dave et al. (4). While CD14 is not itself an IR2 gene, the IR2 signature captures genes preferentially expressed in macrophages and dendritic cells (TLR5, FCGR1A, SEPT10, LGMN, C3AR1), and CD14 marks the same myeloid compartment at the protein level. The intrafollicular niche where CD14-high FDCs and M2 macrophages coexist with tumor B cells may represent the tissue correlate of the IR2 transcriptional program.

The dominance of VISTA over PD-L1 in the FL myeloid compartment has direct therapeutic implications. Current clinical trials in FL focus predominantly on PD-1/PD-L1 axis blockade, which has shown limited efficacy in FL compared to other lymphomas (21). Our data suggest that VISTA, rather than PD-L1, is the operative immune checkpoint in FL, particularly within the myeloid compartment. The association between EP300 mutations and VISTA upregulation on M2 macrophages further suggests that genetic alterations in FL may directly shape the immune evasion landscape, offering a potential avenue for patient stratification.

Integrating these findings, we propose a dual-compartment model of immune evasion in FL (**Figure 7**). Two spatially distinct programs operate on either side of the follicle boundary: an intrafollicular immune-privileged niche organized by CD14-high FDCs and M2 macrophages, and an interfollicular suppressive zone dominated by VISTA+ myeloid cells. A Treg barrier at the follicle-T zone interface separates the two. CD14 integrates these programs as a biomarker, capturing signal from both macrophages and CD14-high FDCs across the follicular-interfollicular axis.

**Figure 7.**
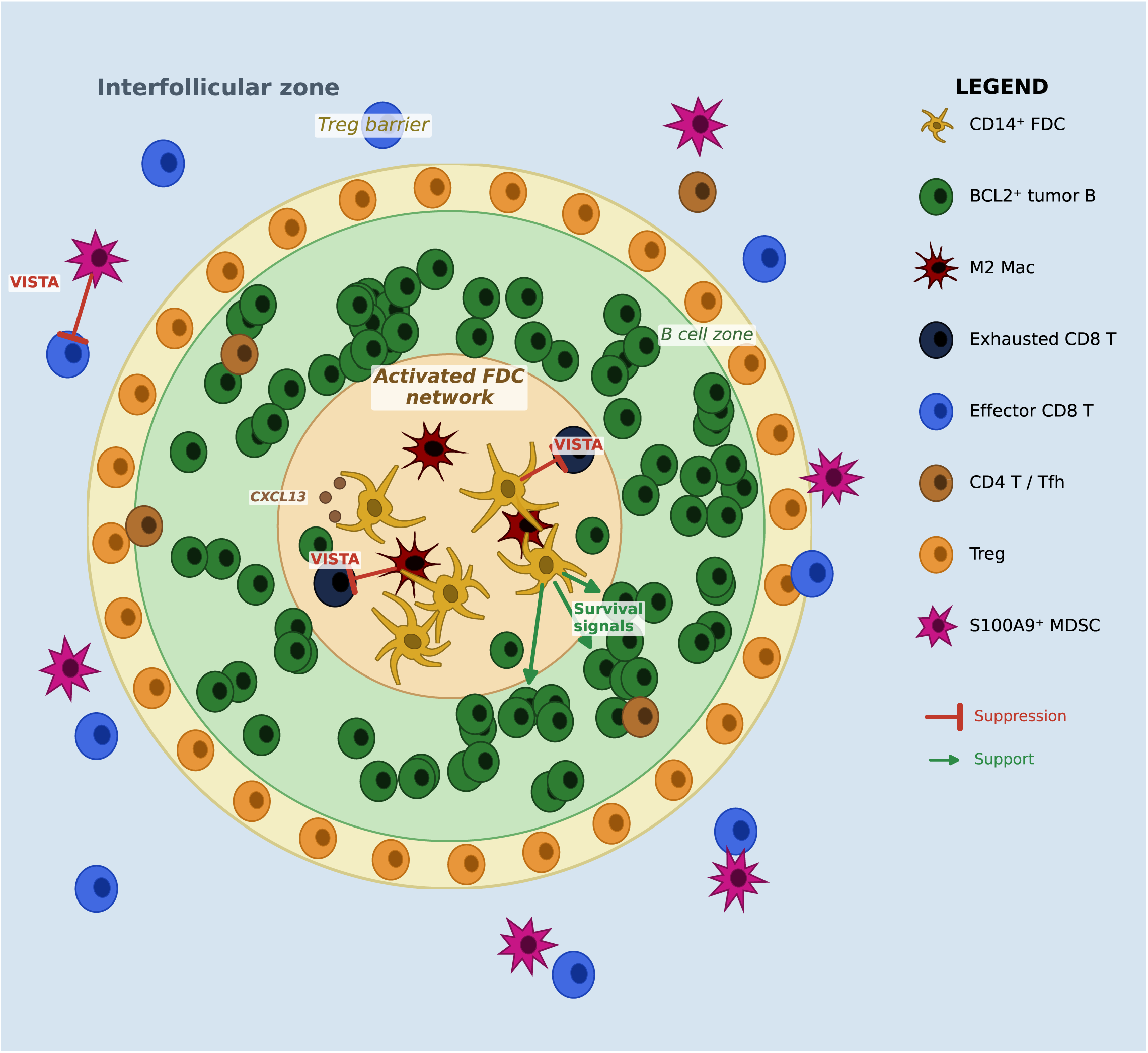
Dual-compartment immune evasion model in follicular lymphoma. Proposed model integrating all spatial and genetic findings. Intrafollicular sanctuary: immune exclusion (2.0% CD8 T, 73.6% exhausted), VISTA+ CD14+ FDCs organize the niche, M2 Mac in FDC zone (41%). Treg barrier at boundary (Treg:CD8 = 1.34) with CD39+ U-shaped gradient. Interfollicular suppression: VISTA+ on 56% of myeloid (vs PD-L1 12%), S100A9+ MDSC-like cells (3.6x in transformers), perivascular niches. Biomarker: CD14 predicts PFS (HR=1.50, p=0.0007) independent of FLIPI, grade, stage, age. EP300 mutation associated with VISTA+ M2 Mac (p=0.003), which associates with transformation (OR=4.48). POD24: CD14+FLIPI AUC=0.77.

Several limitations should be acknowledged. The tissue microarray format captures a 1 mm core per region, which may not represent the full heterogeneity of the lymph node. The cross-panel comparison relies on serial sections rather than simultaneous measurement, though registration analysis confirmed high spatial alignment. Most patients in this cohort were treated with R-CVP, a regimen that has largely been superseded by bendamustine- and rituximab-CHOP-based immunochemotherapy, and the prognostic associations we report will need to be tested under these modern regimens. The CD14 biomarker and the POD24 classifier were derived and evaluated within this single cohort, so their in-sample performance, including the 0.77 area under the curve, will require confirmation in an independent cohort before clinical application. Finally, the agent-based model simplifies the biological complexity and serves primarily as a hypothesis-generating tool rather than a quantitative prediction.

In conclusion, this work establishes that FL exploits tissue architecture as an immune evasion mechanism and identifies CD14-high FDCs and VISTA as actionable targets. The spatial biomarker CD14, already measurable by standard immunohistochemistry as demonstrated by Smeltzer et al. (15), predicts outcomes independently of FLIPI and may enable risk stratification in clinical practice. Therapeutic strategies that breach the follicle boundary, such as bispecific antibodies combined with VISTA blockade, may be required to overcome the architectural immune evasion that defines FL biology.

## METHODS

### Patient cohort and tissue microarrays

Formalin-fixed, paraffin-embedded (FFPE) follicular lymphoma tissue was obtained from three tissue microarrays (TMAs A1, B1, C1) constructed at BC Cancer (Vancouver, Canada) and one commercial TMA (Biomax). Each TMA contained 1 mm diameter cores from diagnostic FL biopsies. Control cores (tonsil, prostate, kidney, spleen, adrenal) were included on each TMA and excluded from tumor analyses. After excluding controls, the dataset comprised 156 tumor cores from 131 patients (139 cores) plus a commercial TMA (17 cores). Clinical data including age, sex, Ann Arbor stage, FLIPI score, histologic grade, treatment history, progression-free survival, overall survival, and transformation status were available for patients on the BC Cancer TMAs.

### Imaging mass cytometry

Serial sections from each TMA were stained with two complementary 39-marker metal-conjugated antibody panels (**Table S1**). The T-panel targeted T cell subsets (CD3, CD4, CD8a, CD45RO, CD127), exhaustion and checkpoint markers (TOX, PD-1, TIM-3, CD39, CTLA-4, LAG-3), B cell markers (CD20, CXCR5), and myeloid markers (CD68). The S-panel targeted myeloid populations (CD14, CD68, CD163, CD206, S100A9, CD11c, CD11b), B cell and FDC markers (CD20, CD21, PAX5, BCL2), stromal markers (Vimentin, PDPN, Fibronectin, SOX9), vascular markers (CD31, CD34, CD146), checkpoint and immune regulation molecules (VISTA, IDO, PD-L1), and chemokines (CXCL13, CXCL12, CCL21). Both panels included structural markers (Histone H3, pH3s28, DNA1, DNA2) for nuclear identification and segmentation. Imaging was performed on a Hyperion Imaging System (Standard BioTools) at 1 µm resolution, producing pixel-level intensity measurements for each marker.

### Cell segmentation

Cells were segmented using a hybrid approach combining nuclear detection with Cellpose boundary prediction. Nuclear channels (DNA1 + DNA2) were smoothed and local maxima detected as seed points. Cellpose (cyto3 model, flow_threshold=0.8) was then applied to predict cell boundaries. Only Cellpose-predicted cells containing at least one detected nucleus were retained, removing approximately 22% of predicted objects (cytoplasm fragments, segmentation artifacts). The elevated flow threshold (0.8 vs default 0.4) was chosen to avoid under-segmentation in dense lymphoma tissue. For each retained cell, mean marker intensities were computed across all pixels within the cell boundary, along with centroid coordinates and cell area.

### Data transformation and quality control

Raw mean intensities were transformed using arcsinh(X/5) with a cofactor of 5, standard for mass cytometry data. Marker quality was assessed by computing the 99th percentile of arcsinh-transformed expression pooled across tumor cores for each TMA. Markers with p99 below 0.3 were classified as low dynamic range; these were noted but not excluded from analysis, as some (VISTA, IDO, CXCL13) retained biological signal despite low overall intensity. Per-marker z-scoring with clipping at 10 was applied for dimensionality reduction and clustering.

### Cell type annotation

Cell types were assigned using hierarchical rule-based gating on arcsinh-transformed marker values rather than clustering, because B cell dominance in FL inflates cluster-mean CD20 expression and prevents reliable identification of minority T cell populations. All 39 markers on each panel were available to the gating rules; the subsets displayed in **Figure 1d,e** are visualization-only summaries of the lineage-defining and functional markers used to define each cell type and are not themselves the basis of the annotation. Gating rules were applied in a priority hierarchy, with each cell assigned to the first matching rule and not re-evaluated against later rules. Cells passing DNA quality control but not meeting any lineage marker threshold were classified as Unassigned. Cells with ambiguous marker profiles that did not fit cleanly into a single lineage (e.g., co-expressing B cell and T cell markers) were classified as Other.

For the T-panel, the gating hierarchy was (in order): (1) CD8 T exhausted: CD3>0.5 AND CD8>CD4 AND CD8>CD20 AND TOX>0.8 AND PD-1>0.5; (2) CD8 T pre-exhausted (TOX+): as above but TOX>0.8 without PD-1 criterion; (3) CD8 T cells: CD3>0.5 AND CD8>CD4 AND CD8>CD20; (4) Treg: CD3>0.5 AND CD4>CD8 AND CD4>CD20 AND FoxP3 above a dynamic threshold (∼1.5× median of positive values, floor 0.3); (5) CD4 T cells: CD3>0.5 AND CD4>CD8 AND CD4>CD20; (6) generic T cells: CD3>0.5 AND (CD4>CD20 OR CD8>CD20); (7) GC B cells (classical CXCR5-high CD20-high germinal-center B cells): CD20>CD3 AND CD20>CD68 AND CD20≥1.5 AND CXCR5>2.0 AND CD20>8.0; (8) Activated B / Plasmablast: CD20>CD3 AND CD20>CD68 AND CD20≥1.5 AND CD38>1.0 AND IRF4>0.5; (9) B cells (CD20hi): CD20>CD3 AND CD20>CD68 AND CD20≥1.5 AND CD20>6.0; (10) B cells (CXCR5hi): CD20>CD3 AND CD20>CD68 AND CD20≥1.5 AND CXCR5>1.5; (11) B cells (TOXhi): CD20>CD3 AND CD20>CD68 AND CD20≥1.5 AND TOX>1.5; (12) B cells (broad class capturing most malignant FL B cells, which typically modulate CXCR5 and CD20 and do not meet the strict GC B cell thresholds): CD20>CD3 AND CD20>CD68 AND CD20≥1.5; (13) Macrophages (GzmB+): CD68>2.0 AND GranzymeB>2.0 (CD68- positive granzyme B-expressing macrophages; earlier iterations labeled these as “Cytotoxic (GzmB+)” but their marker profile is CD68-high and CD3/CD8-negative, consistent with macrophage identity); (16) Macrophages: CD68>2.0 AND CD68>CD3; (17) low-signal relative-CD20 gate: CD20≥0.5 AND CD20>2×CD3 AND CD20>2×CD68 — cells satisfying this relative comparison but lacking any positive lineage marker signal were classified as Unassigned; (18) Mixed / Border: CD20>1.0 AND CD3>0.5. Cells meeting none of the above were classified as Unassigned or Other.

For the S-panel, the gating hierarchy was (in order): (1) FDC: CD21>5.0, or CD21>2.0 AND CXCL13>0.3 AND CD20<CD21; (2) FRC (PDPN+): PDPN>1.5 AND CD20<2.0 AND CD68<2.0; (3) Endothelial: Vimentin>3.0 AND low CD20/CD68/CD4/CD8 AND (CD31>1.5 OR CD34>1.5); (4) Stromal / CAF: Vimentin>3.0 AND low lineage markers; (5) CD4 T cells: CD4>CD20 AND CD4>CD68 AND CD4>0.5; (6) CD8 T cells: CD8>CD20 AND CD8>CD68 AND CD8>0.5; (7) B cells (BCL2+): CD20>1.5 AND CD20>CD68 AND BCL2>2.0; (8) B cells (PAX5+): CD20>1.5 AND CD20>CD68 AND PAX5>1.0; (9) B cells: CD20>1.5 AND CD20>CD68; (10) Myeloid (S100A9+): S100A9>5.0; (11) M2 Macrophages: CD68>2.0 AND (CD163>1.5 OR CD206>1.5); (12) M1 Macrophages: CD68>2.0 AND (CD11c>1.0 OR HLA-DR>2.0); (13) Macrophages: CD68>2.0 OR (CD14>2.0 AND CD14>CD20); (16) Dendritic cells: CD11c>1.5 AND HLA-DR>1.5 AND CD14<1.0; (17) pDC: CD123>1.5; (18) Endothelial: CD31>2.0 OR CD34>1.2; (15) Stromal / CAF: Fibronectin>2.0; (19) Histiocytes (CD44hi): CD44>5.0; (20) Mixed / Border: CD20>1.0 AND CD68>1.0. Cells meeting none of the above were classified as Unassigned or Other.

The T-panel yielded 15 cell types and the S-panel 18 cell types. In dense tissue regions, IMC segmentation can assign signal from adjacent cells to a given cell’s mask, a phenomenon known as segmentation spillover. This is particularly relevant for cell types that are physically enmeshed in neighborhoods dominated by a single lineage: for example, FDCs residing within dense B cell networks showed apparent expression of B cell markers (PAX5, BCL2, CD20) that strongly correlated with the expression levels in their nearest B cell neighbors (Spearman rho=0.57-0.81) and declined with increasing distance to the nearest B cell, whereas the FDC-intrinsic marker CD21 showed no such dependency (rho=0.10). Cell type identity was therefore defined using lineage-specific markers robust to spillover (e.g., CD21 for FDCs, CD3 for T cells), and analyses of marker co-expression were interpreted with this caveat in mind.

### Tissue compartment analysis (UTAG)

Spatial tissue compartments were identified using the Unsupervised Tissue Architecture Graph (UTAG) method (13). Rather than using raw marker intensities as input features, which fails in B cell-dominant tissues due to message-passing homogenization of CD20 signal, we used one-hot encoded cell type labels as input. Each cell’s neighborhood composition within a 50 µm radius was computed, producing a vector encoding the fraction of each cell type among its spatial neighbors. Leiden clustering (resolution 0.5, igraph backend) was applied to the resulting feature matrix, yielding hundreds of raw domains per panel. These were hierarchically merged using cosine distance and Ward linkage on domain composition vectors, and filtered to the biologically interpretable compartments present in multiple patients, yielding nine compartments on the T-panel and six on the S-panel. Compartments were classified as follicular (>50% B cells) or interfollicular based on their dominant cell type composition.

### Cross-panel concordance

Serial sections stained with T- and S-panels were spatially aligned by overlaying the DNA intercalator channels (Iridium 191/193), which label all nuclei identically on both panels regardless of antibody staining. Tissue overlap was assessed visually by pseudo-coloring the S-panel DNA signal in green and the T-panel in magenta; concordant regions appear white. Per-ROI cell type fractions were compared across panels for shared cell populations (CD4 T, CD8 T, B cells, myeloid). Pearson correlations were computed for each cell type across all paired ROIs. Smoothed marker expression maps (Gaussian kernel, sigma=20 µm) were generated for CD20 on both panels to assess spatial concordance at the tissue level.

### Survival analysis

Spatial analyses pooled all cores to maximize statistical power. For survival analyses, one core per patient was used (earliest available timepoint) to ensure independence of observations. Clinical outcomes included progression-free survival (PFS), overall survival (OS), histologic transformation (binary, compared by Mann-Whitney U test), and progression of disease within 24 months (POD24). Univariate Cox proportional hazards models were fitted for each cell type fraction and marker intensity using the lifelines Python package. Hazard ratios were computed per standard deviation increase. Multivariate Cox models for CD14 included progressive covariate adjustment: univariate, then adding FLIPI score, histologic grade, and finally Ann Arbor stage and age. POD24 prediction was assessed using logistic regression with 5-fold stratified cross-validation (repeated with fixed seed for reproducibility); the cross-validated area under the receiver operating characteristic curve (AUC) was computed for CD14 alone, FLIPI alone, and the combined model. Kaplan-Meier curves were generated using median marker intensity splits, with log-rank test P-values. All survival analyses were restricted to treated patients. Of the 131 patients, 14 received observation only and one lacked a diagnostic clinical record (only a relapse biopsy was available), leaving 116 treated patients for these analyses.

### Permutation-based neighborhood enrichment

Pairwise cell-cell interaction enrichment was assessed using a permutation approach. For a random subsample of 60 ROIs (for computational efficiency), K=10 nearest neighbors were identified for each cell within each compartment using a KD-tree. Cell type labels were then permuted 200 times within each ROI to generate a null distribution of neighbor-type counts. Z-scores were computed as (observed - null mean) / null standard deviation. Interactions were filtered to retain only cross-type pairs (excluding self-interactions) with z-score above 5 in at least one compartment and a minimum of 100 observed interactions to prevent spurious enrichment from rare populations.

### Single-cell RNA sequencing re-analysis

Published scRNA-seq data from FL lymph nodes (16) were obtained from CZ CELLxGENE (137,147 cells, 30,172 genes). Analysis focused on FDC (n=108) and myeloid cell populations. CD14 expression was compared between CD14-positive and CD14-negative FDCs. Total UMI counts per cell were compared using the Mann-Whitney U test. Genes correlated with CD14 expression in FDCs were identified by Spearman correlation after library-size correction. Checkpoint molecule expression (VISTA, PD-L1, IDO1) was compared across cell types. All scRNA-seq analyses used Scanpy.

### Agent-based model

An agent-based model was implemented in the Mesa framework (Python) to simulate the dual-compartment immune evasion architecture. The model encoded 8 cell types (tumor B, CD8 T, CD4 T/Tfh, Treg, M1 Mac, M2 Mac, S100A9+ MDSC, FDC), 4 diffusive fields (CXCL13, CXCL9/10, IDO kynurenines, FDC survival signal), and concentric zone geometry (follicle center, B zone, boundary, T zone) on a 100x100 grid. Cell behaviors including proliferation, killing, exhaustion, and suppression were parameterized from published measurements and calibrated against our IMC data. **Table S2** provides a comprehensive parameter justification with literature references for all model parameters. Key parameters included: tumor B cell proliferation rate (0.005/tick, based on Ki-67 positivity rates in FL grade 1-2 (22)), CD8 T cell kill rate (0.08, based on in vivo CTL killing measurements (23)), VISTA-mediated exhaustion (0.008, based on contact-dependent T cell quiescence (24)), and Treg suppressive strength (0.3, based on CTLA-4/IL-10/TGF-beta signaling (25)). Treatment scenarios modeled anti-VISTA (reducing myeloid suppression), rituximab (antibody-dependent killing of CD20+ cells), and bispecific CD20-CD3 (T cell redirection to CD20+ targets). Simulations were run for 2,000 time steps per condition.

### Software and reproducibility

All analyses were performed in Python 3.11 using NumPy, SciPy, Matplotlib, h5py, Scanpy, lifelines, and scikit-learn. Cell segmentation used Cellpose v4 (cyto3 model). UTAG analysis used the published implementation (13). Code is available at https://github.com/oelemento/IMC-FL (analysis) and https://github.com/oelemento/FL-ABM (agent-based model). Processed single-cell data (imaging mass cytometry, both panels) have been deposited at Zenodo under DOI 10.5281/zenodo.20612591.

## Acknowledgments

We thank Dylan McNally for his help with sample processing and antibody panel optimization. O.E. is supported by UL1TR002384, OT2OD032720, P01CA214274 grants from the National Institutes of Health and LLS SCOR grants 180078-02, 7021-20, 180078-01. W.B. was supported by NIH NCI R01 CA270245, LLS TRP 6641-22, LLS TRP 6679-24, FLF CURE FL 224362, LRF FL PRG 226504, ASH Junior Faculty Scholar Award 202414 and 223390, and Gilead Sciences Research Scholars Program GSI 231647-01. AMM is funded by BCU/IFLI RAFL program grant, and BCU SCOR grants 7027-23 and 7029-23.

## CONFLICT OF INTEREST

O.E. is a co-founder and stockholder in Volastra Therapeutics, holds stock in Freenome, serves on the Scientific Advisory Board (SAB) of and holds stock options in Owkin, Harmonic Discovery, and Exai, is an SAB member of Canary Biosciences, and has received research funding from Eli Lilly, J&J/Janssen, Sanofi, AstraZeneca, and Volastra. AMM has consulted for Treeline Biosciences, Ipsen, Regeneron and Abbvie. DWS declares honoraria from Arima Genomics, AstraZeneca, Chugai, Eli Lilly, Kite/Gilead, Roche and Veracyte, and funding from Roche/Genentech. The remaining authors declare no competing interests.

## DECLARATION OF GENERATIVE AI AND AI-ASSISTED TECHNOLOGIES

During the preparation of this work the authors used Claude (Anthropic) to assist with manuscript drafting, analysis code, and figure preparation. The authors reviewed and edited the content as needed and take full responsibility for the content of the published article.

---

**Figure S1.**
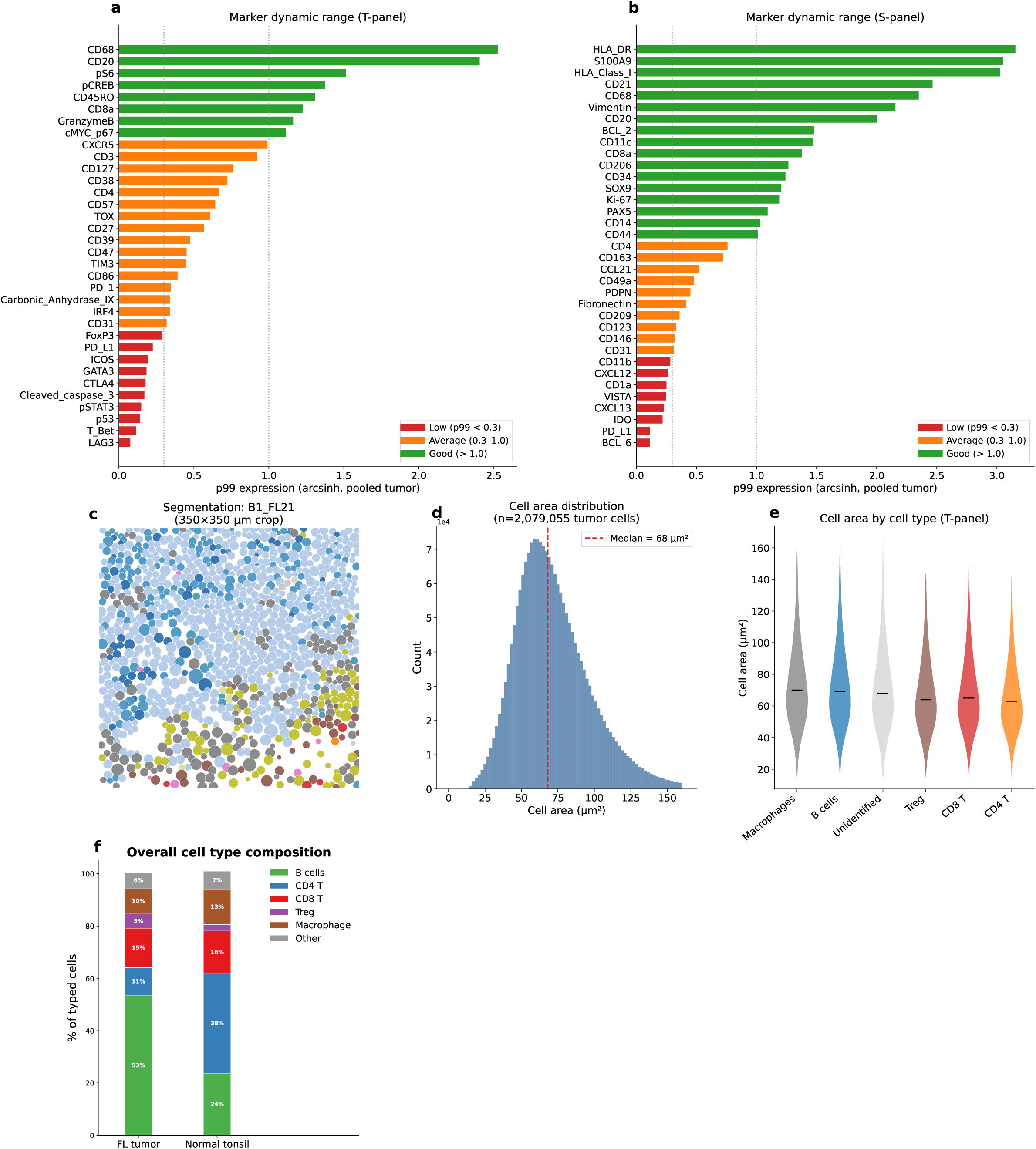
Segmentation and marker quality control. (a) T-panel marker dynamic range (p99 arcsinh expression, pooled tumor cores): green = good (>1.0), orange = average (0.3–1.0), red = low (<0.3). Low-dynamic-range markers: LAG3, T-Bet, CTLA4, p53, pSTAT3, PD-L1, FoxP3, ICOS, GATA3. (b) S-panel marker dynamic range: low-dynamic-range markers include BCL6, PD-L1. (c) Cellpose hybrid segmentation illustration (350×350 µm crop). (d) Cell area distribution (T-panel, median 68 µm²). (e) Cell area by cell type (T-panel violin). (f) Overall cell type composition: FL tumor vs normal tonsil (stacked bar).

**Figure S2.**
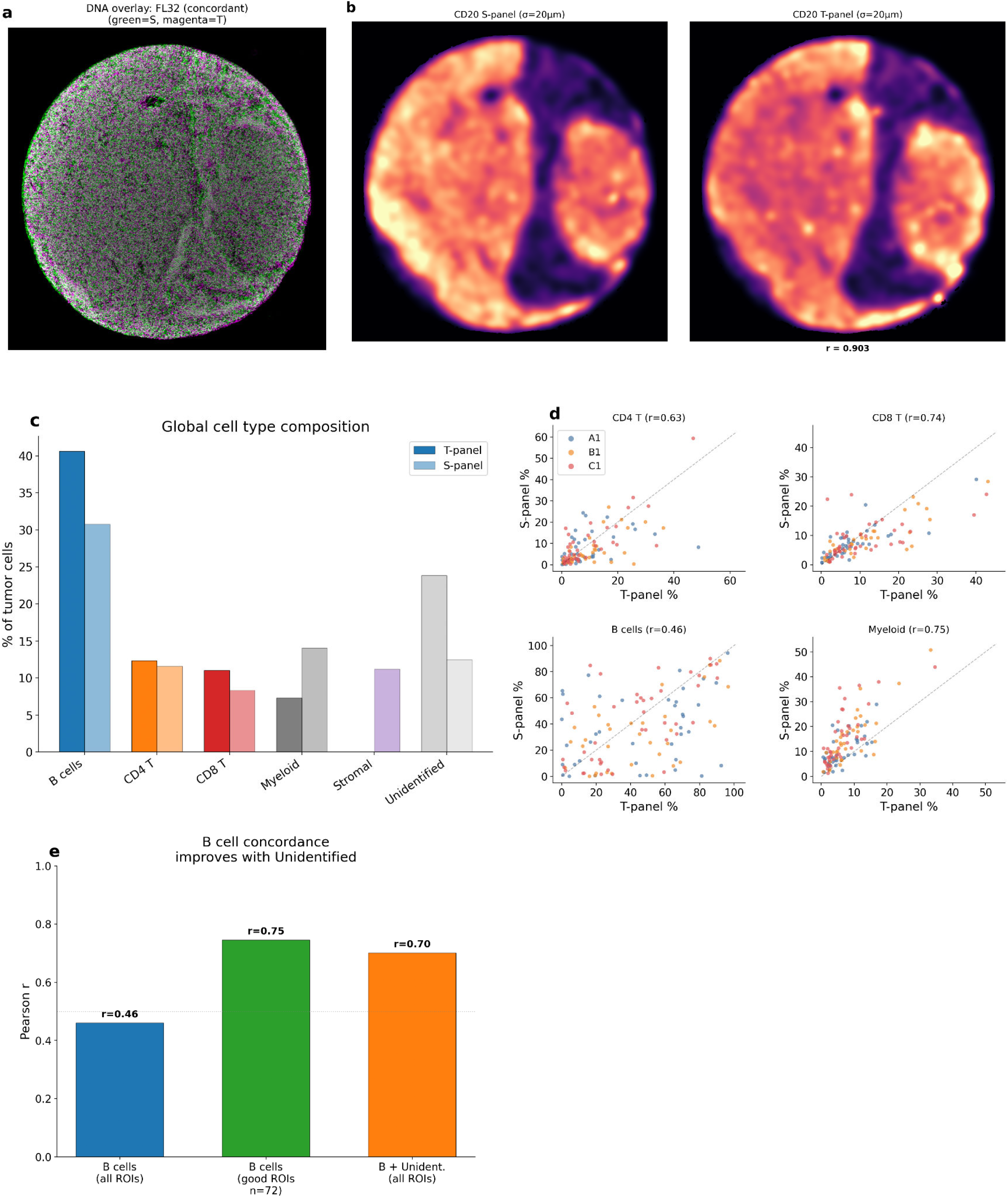
Cross-panel concordance: T-panel vs S-panel on serial sections. (a) DNA-based serial-section registration: fluorescence overlay (green=S, magenta=T) confirms high spatial alignment (FL32 concordant example). (b) Smoothed CD20 expression maps from S-panel and T-panel on the same tissue section (r=0.903). (c) Global cell type composition in both panels (broad categories). (d) Per-ROI scatter: T-panel vs S-panel fractions for CD4 T, CD8 T, B cells, Myeloid (n=115 paired ROIs; Pearson r=0.63-0.75 for CD4 T, CD8 T, and Myeloid; r=0.46 for B cells). (e) B cell concordance improves from r=0.46 (all ROIs) to r=0.75 (good ROIs) when excluding low-quality ROIs.

**Figure S3.**
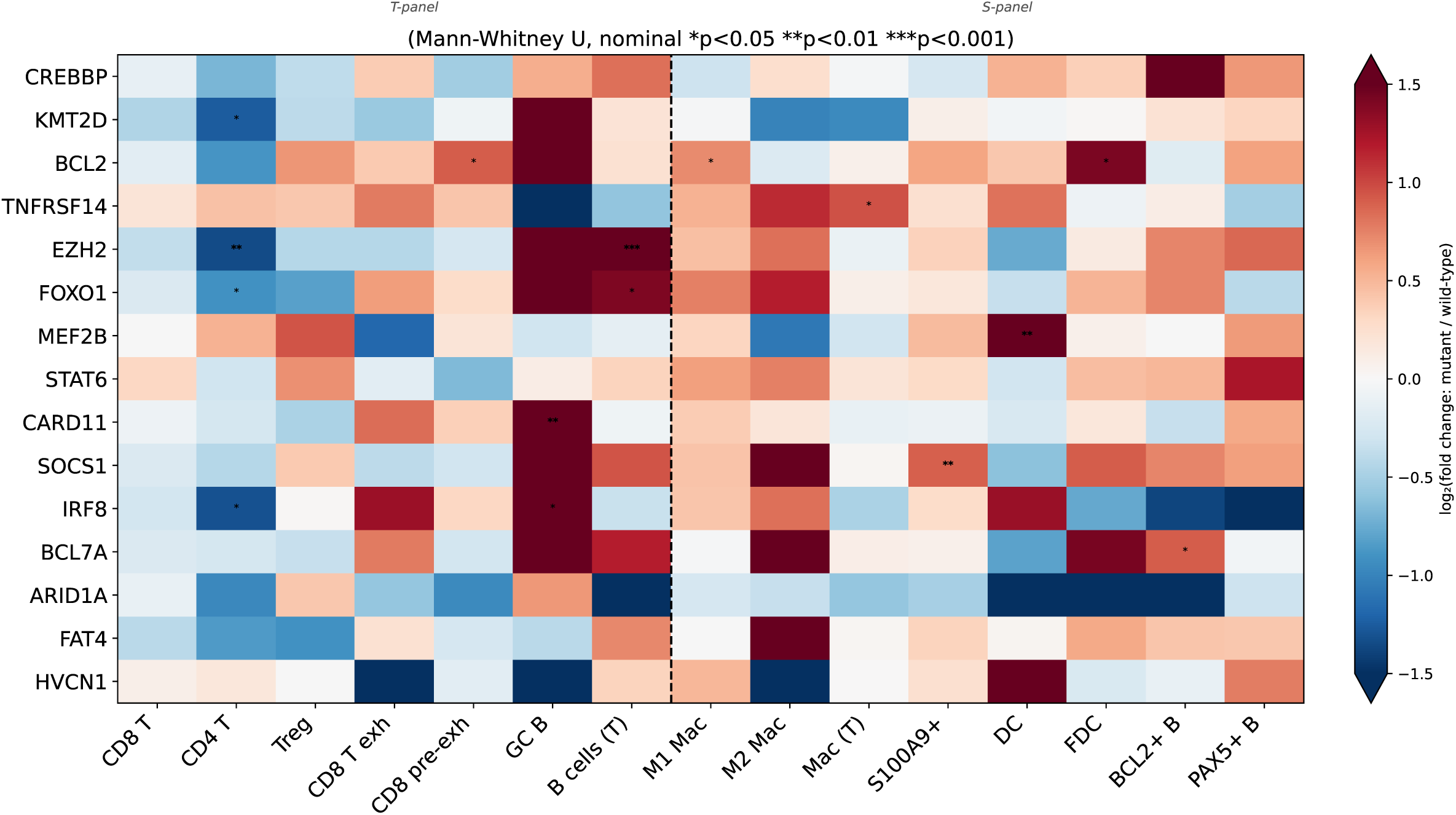
Mutation × cell type fraction associations. Heatmap of log₂ fold change in cell type fractions between mutant and wild-type patients for 15 recurrently mutated genes. Stars indicate nominal significance (*p<0.05, p<0.01,* **p<0.001, Mann-Whitney U). None of 555 tests survive FDR correction (all q>0.07). Mutations in FL do not have detectable large-scale effects on cell type composition as measured by IMC.

**Figure S4.**
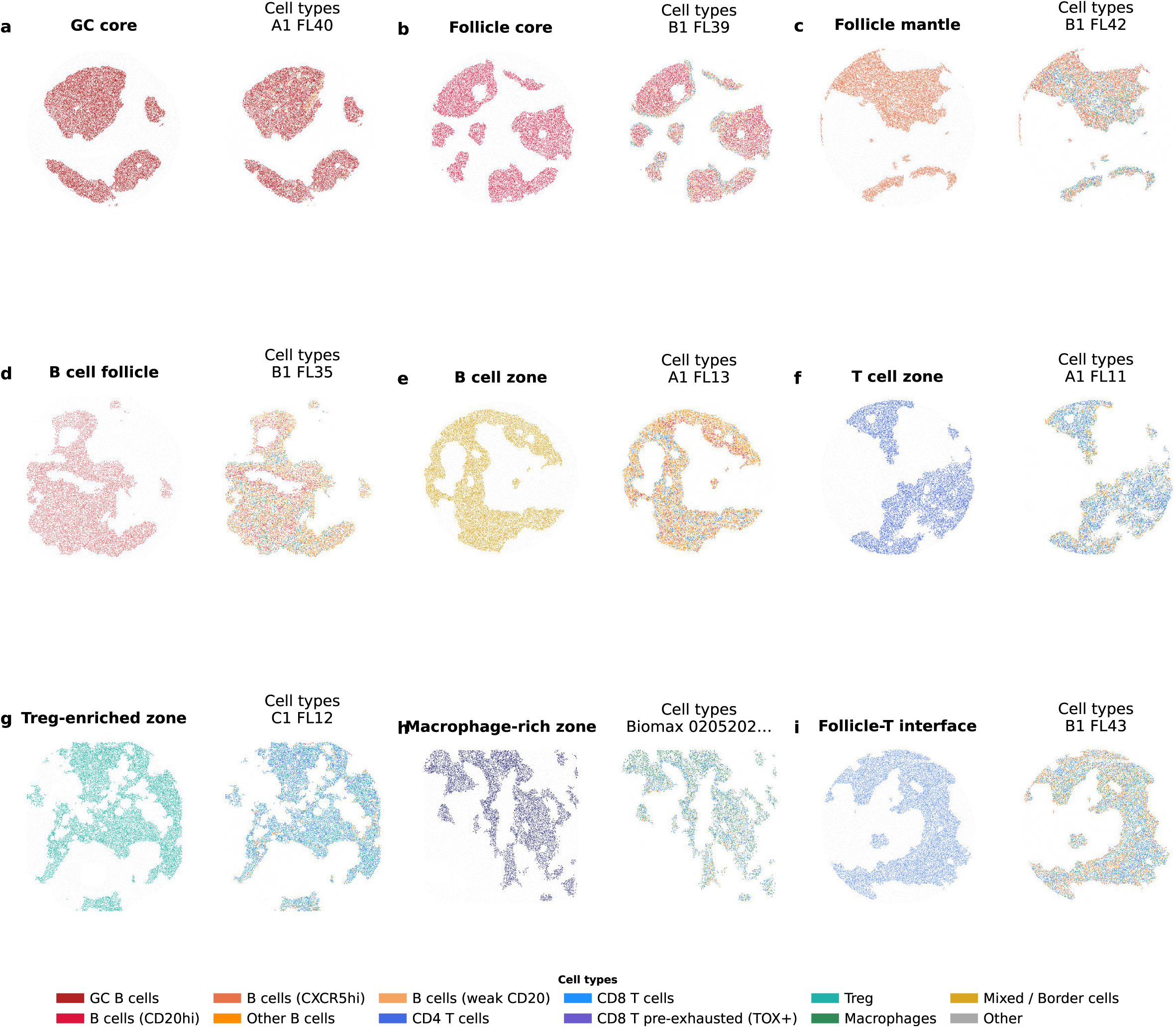
Compartment spatial examples across representative ROIs. Gallery of 9 key T-panel compartments. Each pair shows the compartment highlighted in a representative ROI (left) and cell types within that compartment only (right). Follicular: GC core, Follicle core, Follicle mantle, B cell follicle, B cell zone. Interfollicular: T cell zone, Treg-enriched zone, Macrophage-rich zone, Follicle-T zone interface.

**Figure S5.**
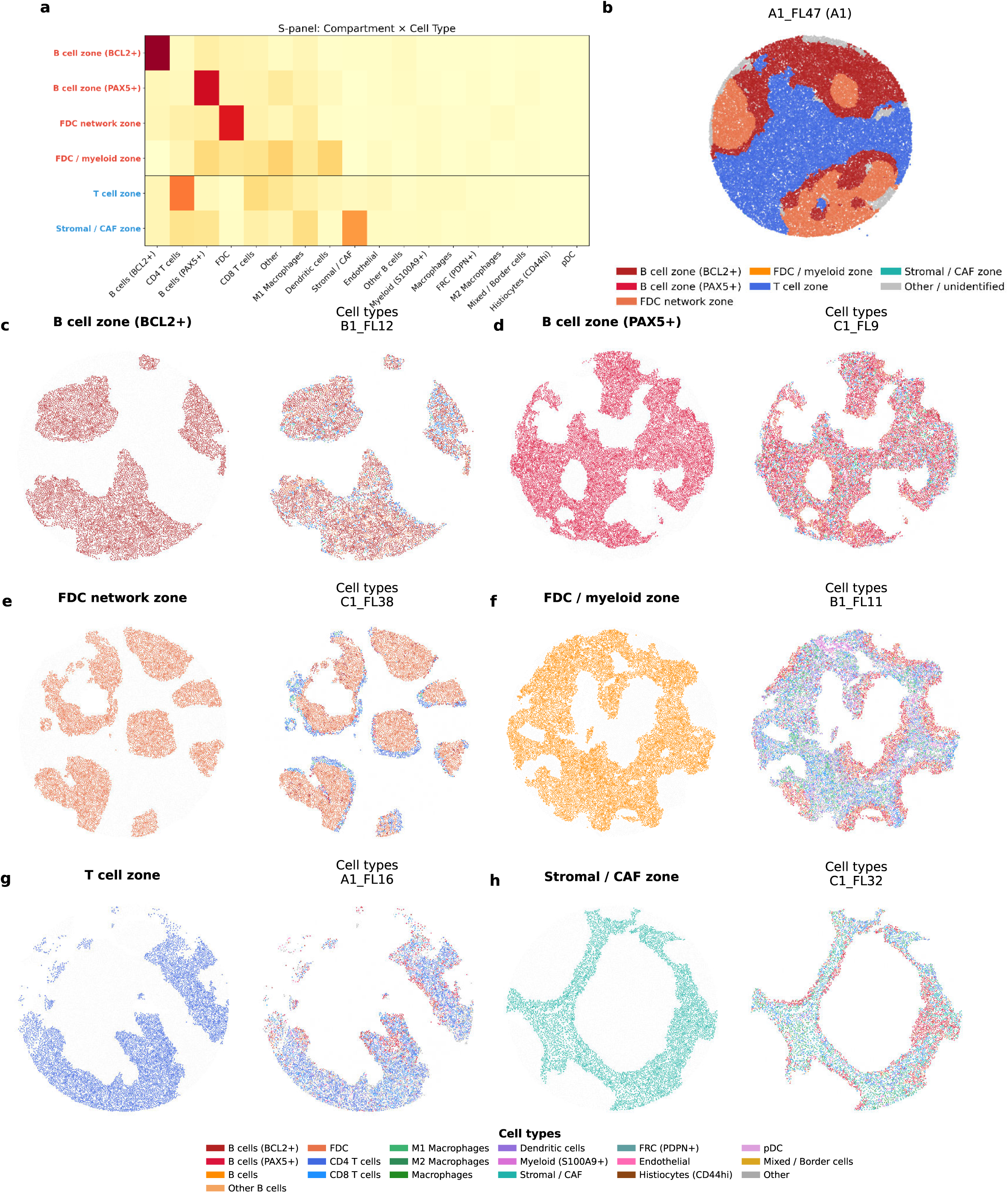
S-panel spatial tissue compartments. S-panel UTAG tissue compartments. (a) S-panel cell type composition per compartment (heatmap). (b) Representative ROI colored by compartment. (c-h) Six key compartments: each pair shows compartment highlighted (left) and cell types within that compartment only (right). Follicular compartments: B cell zone (BCL2+), B cell zone (PAX5+), FDC network zone, FDC/myeloid zone. Interfollicular: T cell zone, Stromal/CAF zone.

**Figure S6.**
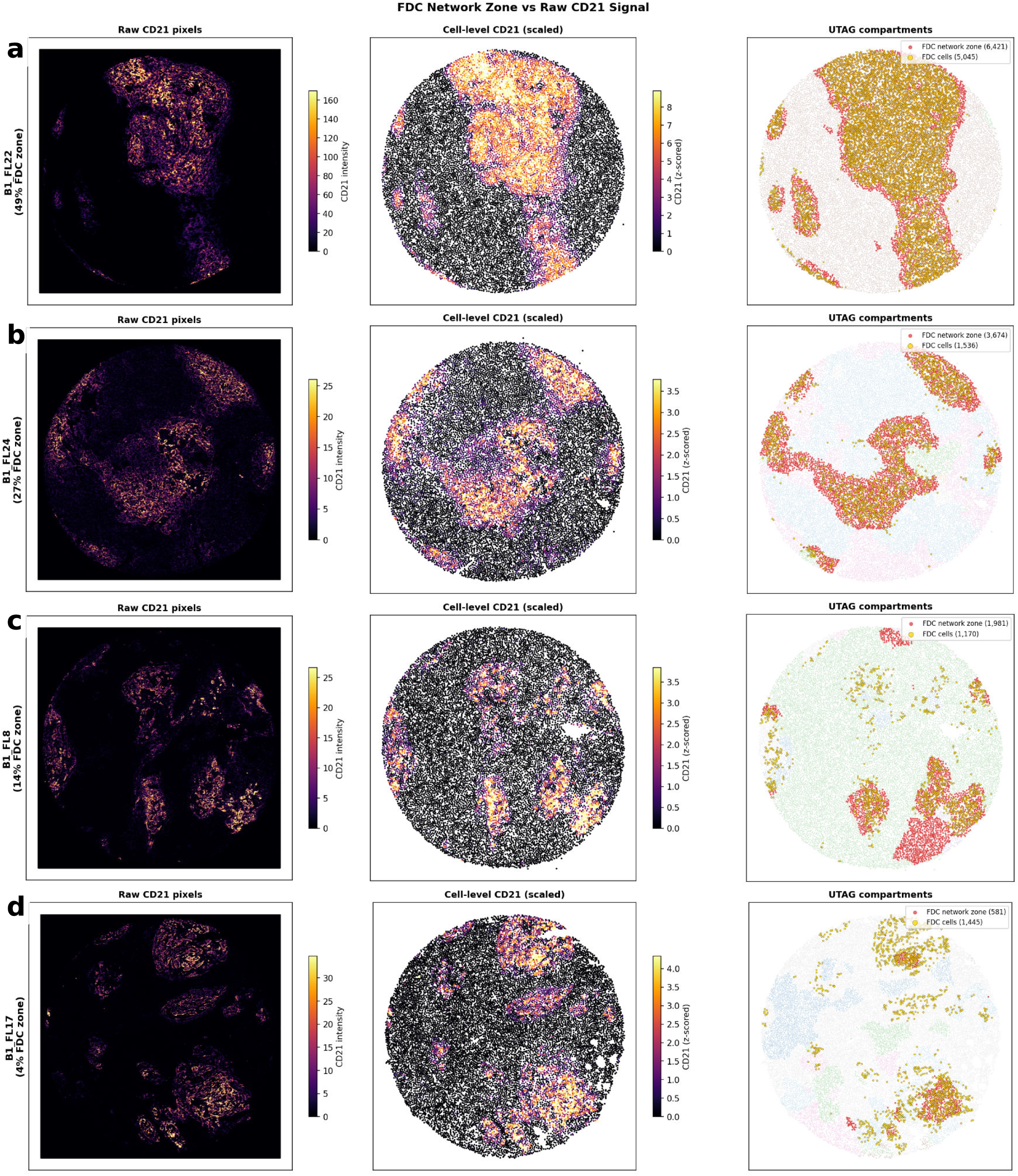
FDC network zone validation: concordance with raw CD21 signal. Spatial concordance between UTAG-defined FDC network zone and raw CD21 IMC signal across 4 B1 ROIs with decreasing FDC zone fraction (49%, 27%, 14%, 4%). Left: raw CD21 pixel intensities (inferno colormap, per-ROI 99.5th percentile clip). Center: cell-level CD21 expression (z-scored, after segmentation and transformation). Right: UTAG compartment map with FDC network zone highlighted in red and individual FDC cells as gold dots. The FDC network zone tracks dense CD21 meshworks faithfully.

**Figure S7.**
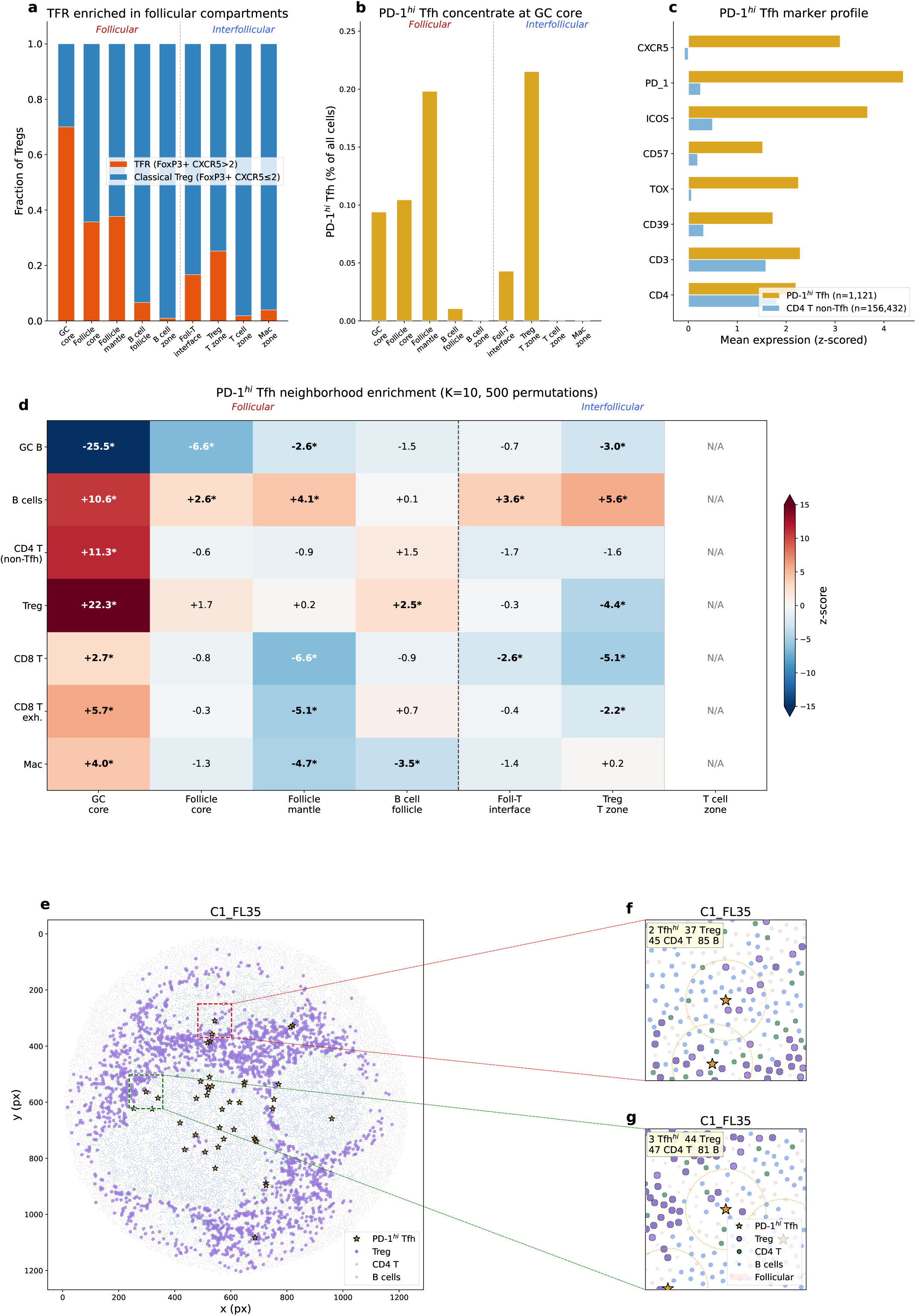
TFR composition, PD-1-hi Tfh niches, and spatial co-localization. (a) TFR (FoxP3+CXCR5>2) vs classical Treg composition across compartments: TFR fraction increases from <2% in T cell zones to 70% at the GC core, indicating that intrafollicular Tregs are predominantly T follicular regulatory cells. (b) PD-1-hi (>1.5) Tfh density (% of all cells) by compartment: highest at GC core. PD-1-lo Tfh at boundary compartments likely include Tfr and CXCR5 spillover artifacts. Tfh are rare (median 3/ROI, 0.03% of typed cells) and do not predict survival (PFS rho=-0.07, p=0.46), transformation (p=0.61), or EZH2 status (p=0.59). (c) Marker profile: PD-1-hi Tfh vs CD4 non-Tfh — PD-1-hi Tfh show elevated CXCR5, PD-1, TOX, and FOXP3 compared to bulk CD4 T cells. (d) Permutation-based neighborhood enrichment (K=10, 500 permutations): PD-1-hi Tfh at GC core are enriched for Tregs (z=+22.3), B cells (z=+10.6), and CD4 T cells (z=+11.3), forming a structured multi-cell hub. (e) Full ROI scatter with PD-1-hi Tfh (gold stars), Tregs (purple), B cells (blue), CD4 T (green); zoom rectangle indicates region shown in (f). (f-g) Two zoom examples of GC-core Tfh-Treg-B cell hubs, showing spatial co-localization within follicular domains (red tint).

**Figure S8.**
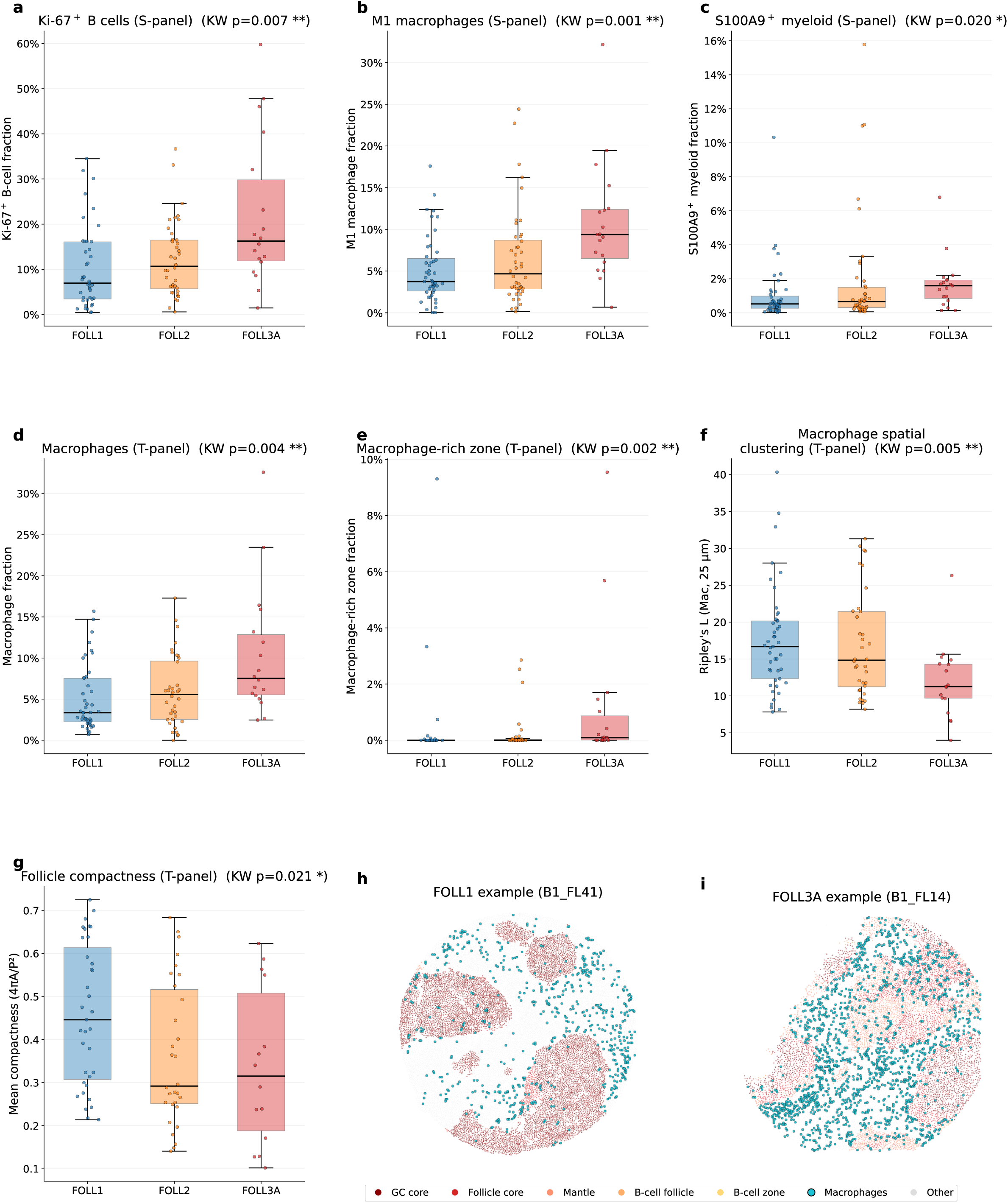
Grade-associated changes in proliferation, myeloid composition, and follicle architecture. Nine-panel supplementary figure (3×3 grid). Patient-level metrics (ROIs averaged per patient before testing) stratified by centrally reviewed histologic grade (FOLL1, FOLL2, FOLL3A). Box plots: black line = median, box = IQR, whiskers = 1.5×IQR (no outliers shown); overlaid points are individual patients. P-values from Kruskal–Wallis across the three grades; significance markers in title: *p<0.05, p<0.01,* **p<0.001. (a) Ki-67⁺ B-cell fraction (S-panel; B cells defined as B cells, B cells (BCL2+), B cells (PAX5+); Ki-67 positive at arcsinh > 0.5; ROIs with ≥200 B cells and ≥8000 typed cells). Included as an internal consistency check that the IMC-derived single-cell B-cell typing recovers the centroblast-count gradient that defines histologic grade. (b) M1 Macrophage cell-type fraction (S-panel). (c) S100A9⁺ Myeloid (MDSC-like) cell-type fraction (S-panel). (d) Macrophage cell-type fraction (T-panel; combined Macrophages and Macrophages (GzmB+)). (e) Macrophage-rich-zone UTAG compartment fraction (T-panel; the compartment is absent in FOLL1 and FOLL2 and emerges in FOLL3A). (f) Macrophage spatial clustering via Besag’s L at radius r=25 µm (T-panel; L(r) = √(K(r)/π) − r, where K is Ripley’s K function; L = 0 under complete spatial randomness, L > 0 indicates clustering, L < 0 indicates dispersion). Lower values at FOLL3A indicate that macrophages occupy a tissue-wide rather than niche-confined distribution. (g) Mean follicle compactness (T-panel). Compactness is the isoperimetric ratio 4πA/P² where A and P are the area and perimeter of each connected follicular domain on a 25 µm binary mask (cells in any of the 5 T-panel follicular compartments: GC core, Follicle core, Follicle mantle, B cell follicle, B cell zone), after morphological closing-then-opening; follicles ≥50 grid pixels (≈31,000 µm²) are kept and the size-weighted mean is reported per ROI. A perfect Euclidean circle scores 1.0; deviation from circularity (elongation, fingering, jagged edges) decreases the value. On a discrete 25 µm pixel mask, the perimeter slightly overestimates the true perimeter so a digital circle scores ≈0.7 in practice; values should be interpreted relatively across grades rather than as a normalized [0, 1] score. (h) Representative FOLL1 ROI (B1_FL41): three discrete circular follicular domains separated by interfollicular tissue, with sparse macrophages (cyan) clustered in the interfollicular spaces. ROI selected by composite ranking across compactness, Ripley clustering, and macrophage fraction, restricted to ROIs with ≥2 distinct follicles and metrics above the FOLL1 median for compactness and below the median for macrophage fraction. (i) Representative FOLL3A ROI (B1_FL14): single irregular follicular mass with dense, dispersed macrophage infiltration throughout. Color key (below panels h and i): follicular sub-compartments shaded from GC core (dark red) through mantle (salmon) to B-cell zone (light yellow); macrophages cyan with black edge; all other cells light gray. Sample sizes per panel after QC filters: panels (a–c, S-panel) n=110 patients (47 FOLL1, 44 FOLL2, 19 FOLL3A); panels (d–g, T-panel) n=99 patients (45 FOLL1, 36 FOLL2, 18 FOLL3A) with additional ≥50 macrophage and ≥200 follicular-cell thresholds for (f) and (g) reducing n further to n=85 and n=79 respectively. None of the per-metric p-values include cross-metric multiple testing correction within the figure; the architectural metrics in (f, g) are interpreted in conjunction with the cell-fraction metrics in (a–e) as a single coherent grade signature rather than as independent tests.

**Figure S9.**
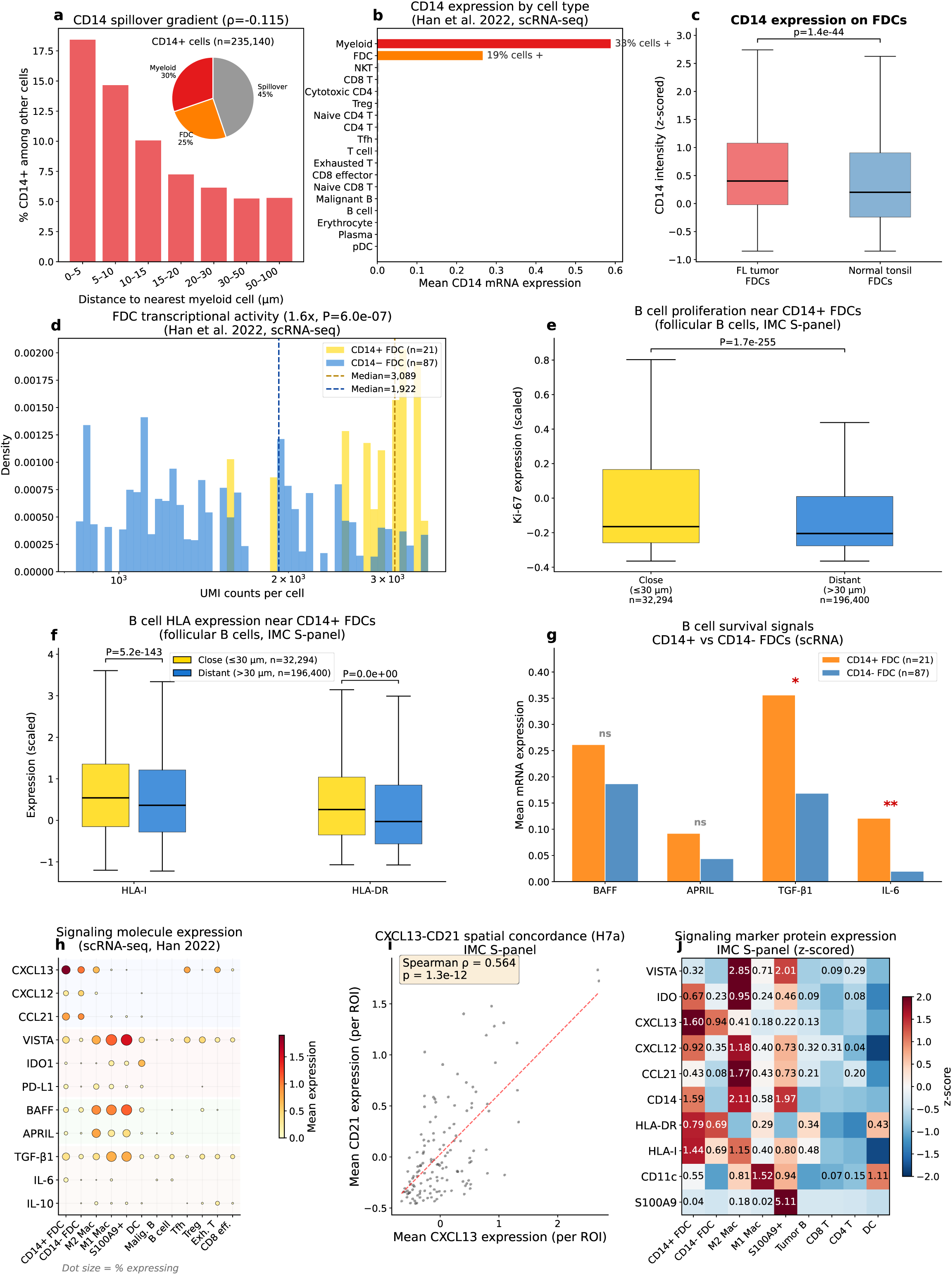
CD14+ FDC validation and signaling characterization. (a) CD14+ cell composition (pie: myeloid 30%, FDC 25%, spillover 45%) and distance-dependent spillover gradient (rho=-0.115). (b) scRNA-seq validation (Han 2022): CD14 mRNA highest in myeloid, second in FDC (19% FDCs positive), confirming IMC protein finding. (c) CD14 expression on FDCs: FL tumor vs normal tonsil (Mann-Whitney); FL FDCs have significantly elevated CD14 (p=1.4e-44). (d) FDC transcriptional activity: CD14+ FDCs have 1.6x higher UMI counts per cell (median 3,009 vs 1,922, P=6.0e-7), confirming CD14 marks a transcriptionally hyperactive state rather than myeloid lineage. (e) B cell proliferation: follicular B cells within 30 px of CD14-high FDCs have higher Ki-67 (mean=0.483 vs 0.301, P=2.9e-32). (f) B cell HLA expression near CD14+ FDCs. (g) B cell survival signals (BAFF, APRIL, TGF-β1, IL-6) in CD14+ vs CD14-FDCs; TGF-β1 (*) and IL-6 (*\*) significantly elevated; APRIL trending (p=0.13), BAFF unchanged. (h) Signaling molecule expression across cell types (scRNA-seq dot plot): FDCs are the dominant source of CXCL13 and CCL21; BAFF and APRIL are predominantly myeloid-derived; myeloid cells also dominate VISTA. (i) CXCL13-CD21 per-ROI concordance (ρ=0.564, p=1.3e-12), confirming FDC-derived CXCL13 organizes the follicular niche. (j) IMC protein heatmap (z-scored): 10 signaling markers × 8 cell types; VISTA peaks on M2 Mac, CXCL13 on FDC, CD14/S100A9 on MDSC-like cells.

**Figure S10.**
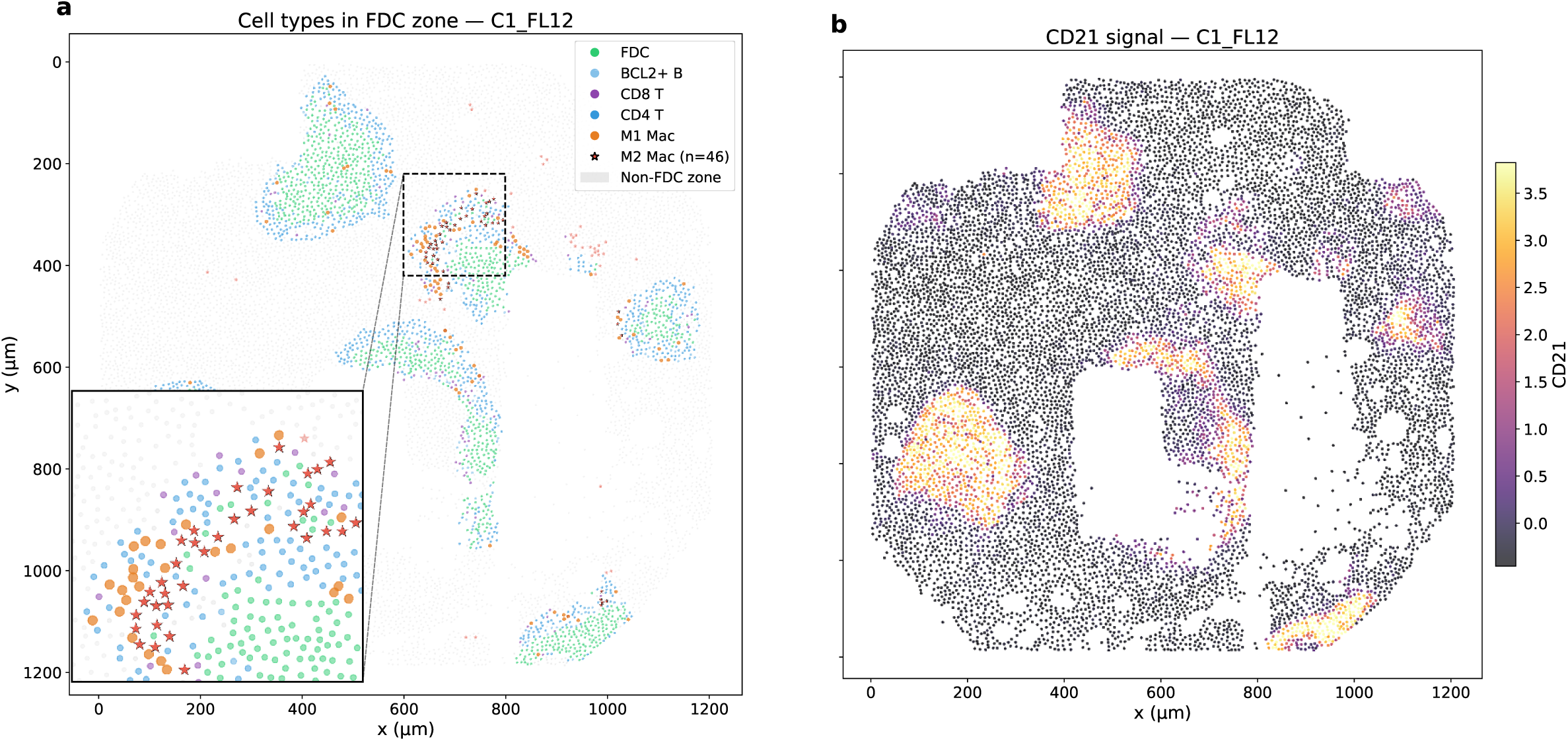
M2 Macrophages in FDC network zone: representative ROI with CD21 validation. (a) Representative ROI (C1_FL12, 46 M2 in FDC zone). Left: cell types in FDC network zone with M2 Mac as red stars; inset zooms on the densest M2 niche. Non-FDC zone cells shown in gray. Right: CD21 signal (inferno colormap) validates that UTAG-defined FDC network zone corresponds to CD21-bright FDC meshwork. M2 Macs populate the FDC network zone but remain spatially separated from CD21-bright FDCs at the single-cell level, consistent with the neighborhood depletion quantified in Fig 5d.

**Figure S11.**
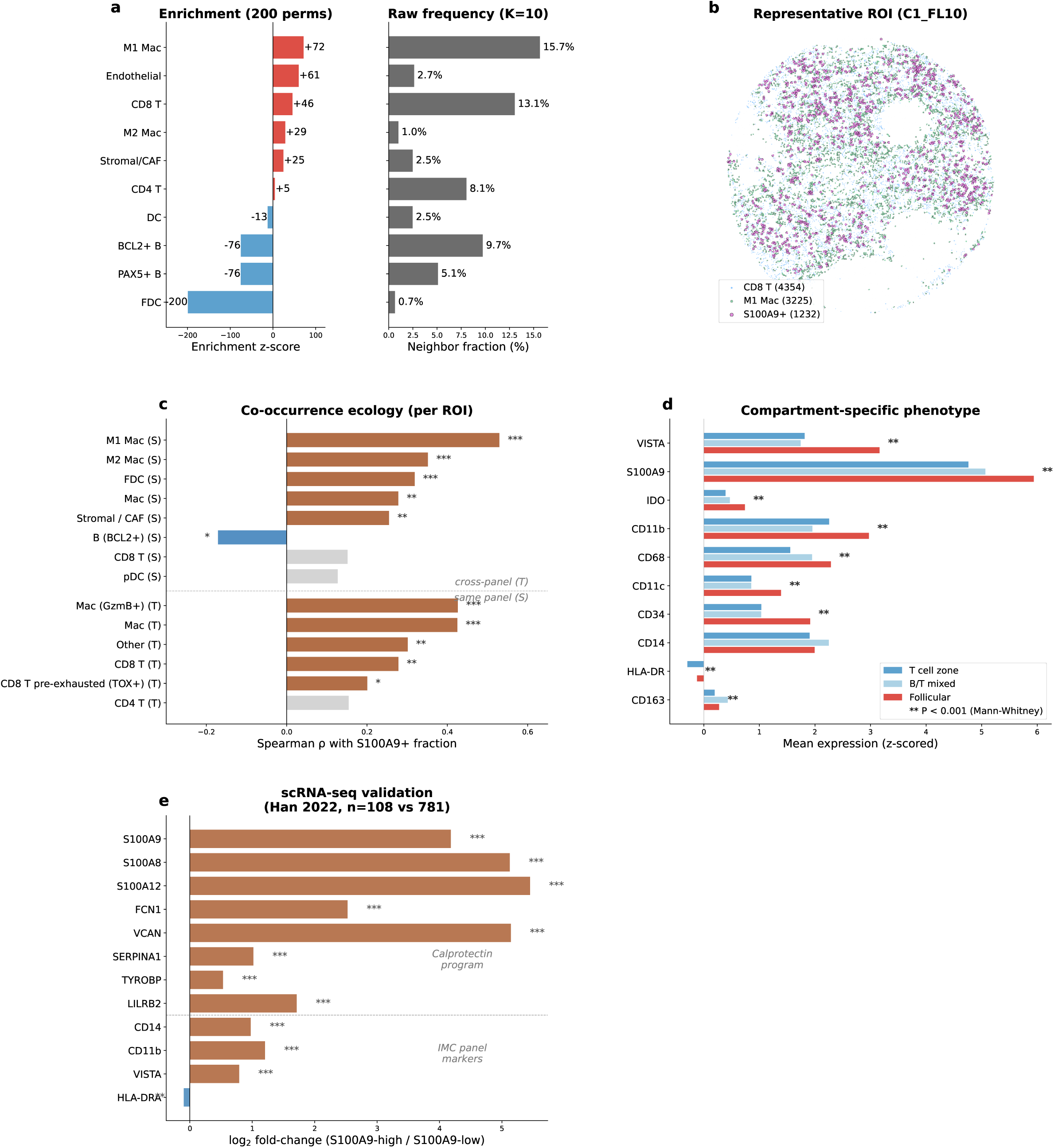
S100A9+ MDSC-like myeloid cells in follicular lymphoma. (a) Neighborhood enrichment z-scores (K=10, 200 permutations): S100A9+ cells are strongly enriched near M1 macrophages (z=+72), endothelial cells (z=+61), and CD8 T cells (z=+46), while depleted near B cells. (b) Representative ROI (C1_FL10): S100A9+ (brown), M1 Mac (red), CD8 T (gold). (c) Co-occurrence ecology: per-ROI Spearman correlation of S100A9+ fraction with other cell types — S100A9+ co-occurs with M1 Mac (ρ=+0.53), GzmB+ macrophages (ρ=+0.43 cross-panel), and FDC (+0.32), marking inflamed microenvironments. (d) Compartment-specific phenotype: follicular S100A9+ express higher VISTA (3.17 vs 1.82), CD68, and CD11b than interfollicular, consistent with an elevated suppressive phenotype; CD14 unchanged across compartments. (e) scRNA-seq validation (Han 2022, 108 vs 781 myeloid cells): calprotectin program (S100A8/A9/A12), FCN1, VCAN, TYROBP upregulated; CD14, CD11b, VISTA concordant with IMC; HLA-DRA unchanged.

**Figure S12.**
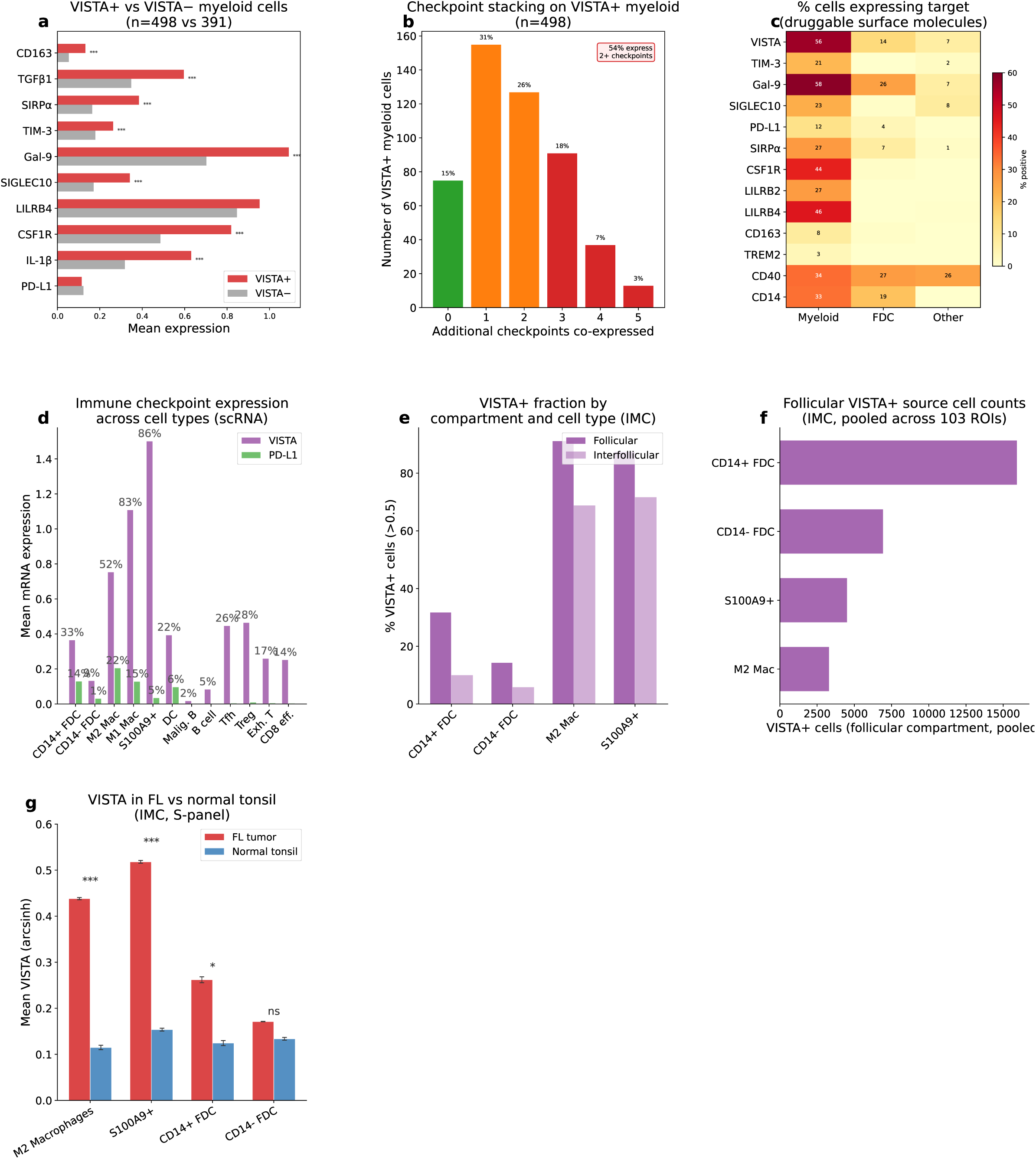
VISTA targeting landscape and checkpoint signaling. (a) VISTA+ vs VISTA− myeloid gene expression (scRNA-seq): VISTA+ cells co-express CD163, TGF-β1, TIM-3, Gal-9, confirming M2-skewed suppressive phenotype. (b) Checkpoint stacking: 94% of VISTA+ myeloid co-express ≥2 additional checkpoints (PD-L1, TIM-3, IDO1, SIGLEC10, Gal-9, PD-L2). (c) Druggable target heatmap: % cells expressing surface molecules across myeloid, FDC, and other cell types. (d) Checkpoint landscape at the transcript level (scRNA-seq, Han 2022): VISTA (VSIR) dominates PD-L1 (CD274) in myeloid cells (56% vs 12% positive), confirming that VISTA — not PD-L1 — is the operative myeloid checkpoint in FL. (e) VISTA+ fractions (% cells > 0.5 scaled) by compartment and cell type (IMC). Per-cell hierarchy in follicle: M2 Mac 91%, S100A9+ 88%, CD14+ FDC 32%, CD14- FDC 15%; all populations substantially reduced in interfollicular compartments. (f) Absolute VISTA+ cell counts in the follicular compartment (pooled across tumor cores, IMC): CD14+ FDCs are the largest VISTA+ source population in the follicle, reflecting their high abundance. (g) VISTA in FL vs normal tonsil (IMC, S-panel). Bars = pooled per-cell mean ± SEM across cells; stars from per-ROI Mann-Whitney (one-sided, FL > tonsil): *p<0.05, p<0.01,* **p<0.001. Per-cell fold change FL/tonsil: M2 Mac 3.8x, S100A9+ 3.4x, CD14+ FDC 2.1x, CD14- FDC 1.3x (ns). See also **Figure S9**(e-h) for additional signaling characterization.

**Table S1.** Imaging mass cytometry antibody panels. Metal-conjugated antibody panels used for imaging mass cytometry. (a) T-panel (T/B/NK, immune-focused), targeting T cell subsets, exhaustion and checkpoint markers, B cell markers, and myeloid markers. (b) S-panel (stromal/myeloid-focused), targeting myeloid populations, B cell and follicular dendritic cell markers, stromal and vascular cells, and immune checkpoint and chemokine molecules. Each row lists the metal isotope tag, antibody target, working dilution, clone, vendor, catalog number, and lot number. DNA intercalators (Ir191, Ir193) used for nuclear identification and segmentation are not antibody-based and are not listed.

| Metal ID | Antibody | Dilution | clone | Vendor | Catalog ID | Lot ID |
| --- | --- | --- | --- | --- | --- | --- |
| 113In | total H3 | 1:400 | D1H2 | CST | 4499BF | 17 |
| 115In | p-H3 s28 | 1:100 | HTA28 | BioLegend | 641002 | B241679 |
| 141Pr | CD38 | 1:50 | EPR4106 | abcam | ab226034 | GR3347999-1 |
| 142Nd | Cleaved Caspase 3 | 1:50 | 5A1E | CST | 9664BF | 10 |
| 143Nd | CXCR5 | 1:50 | D6L3C | CST | 72172BF | 3 |
| 144Nd | CD57 | 1:100 | HNK-1 | BioLegend | 359602 | B287684 |
| 145Nd | TIM3 | 1:50 | D5D5R | CST | 45208BF | 8 |
| 146Nd | PD-L1 | 1:25 | SP142 | abcam | ab236238 | GR3246745-8 |
| 147Sm | CD278 (ICOS) | 1:100 | D1K2T | CST | 89601BF | 6 |
| 148Nd | H3K27me3 | 1:200 | C36B11 | CST | 9733BF | 17 |
| 149Sm | T bet | 1:50 | D6N8B | CST | 13232BF | 5 |
| 150Nd | PD-1 | 1:50 | D4W2J | CST | 86163BF | 7 |
| 151Eu | CD31 | 1:50 | EPR3094 | abcam | ab207090 | GR3229164-6 |
| 152Sm | CD86 | 1:100 | E2G8P | CST | 91882BF | 2 |
| 153Eu | LAG3 | 1:50 | D2G40 | CST | 15372BF | 5 |
| 154Sm | Tox | 1:100 | NAN448B | abcam | ab256486 | GR3345326-1 |
| 155Gd | FoxP3 | 1:25 | 236A/E7 | Invitrogen | 14-4777-82 | 2129676 |
| 156Gd | CD4 | 1:50 | EPR6855 | abcam | ab181724 | GR3285644-12 |
| 158Gd | p-stat3 | 1:50 | D3A7 | CST | 9145BF | 39 |
| 159Tb | CD68 | 1:100 | KP1 | Invitrogen | 14-0688-82 | 24B4431 |
| 160Gd | CD39 | 1:50 | EPR20627 | abcam | ab236038 | GR3274485-5 |
| 161Dy | CD20 | 1:100 | L26 | Invitrogen | 14-0202-82 | 2380076 |
| 162Dy | CD8a | 1:100 | C8/144B | Invitrogen | 14-0085-82 | 2151435 |
| 163Dy | CD27 | 1:25 | EPR8569 | abcam | ab256583 | GR3349592-2 |
| 164Dy | c-Myc p67 | 1:50 | 9E10 | SBT | 3164025D | 0651905 |
| 165Ho | p-CREB | 1:50 | 87G3 | SBT | 3165034D | 2851605 |
| 166Er | GATA3 | 1:25 | L50-823 | BD | 558686 | 9346413 |
| 167Er | Granzyme B | 1:50 | EPR20129-217 | abcam | ab219803 | GR3349314-3 |
| 168Er | CD127 | 1:50 | EPR2955(2) | abcam | ab240225 | GR3375511-4 |
| 169Tm | CD47 | 1:50 | EPR21794 | abcam | ab233122 | GR3217258-4 |
| 170Er | CD3 | 1:100 | Polyclonal | SBT | 3170019D | 220222-16 |
| 171Yb | CTLA4 | 1:50 | CAL49 | abcam | ab251599 | GR3274686-11 |
| 172Yb | IRF4 | 1:50 | EP5699 | abcam | ab240071 | GR3255392-1 |
| 173Yb | CD45 RO | 1:400 | UCHL1 | BioLegend | 304202 | B265382 |
| 174Yb | Carbonic Anhydrase IX | 1:200 | Polyclonal | R&D systems | AF2188 | VNQ0319011 |
| 175Lu | p-S6 | 1:50 | D57.2.2E | CST | 4858BF | 20 |
| 176Yb | p53 | 1:50 | 7F5 | CST | 2527BF | 11 |

Table S1. Antibody Panel: S-panel (stromal/myeloid)
| Metal ID | Antibody | Dilution | clone | Vendor | Catalog ID | Lot ID |
| --- | --- | --- | --- | --- | --- | --- |
| 113In | total H3 | 1:400 | D1H2 | CST | 4499BF | 17 |
| 115In | p-H3 s28 | 1:100 | HTA28 | BioLegend | 641002 | B241679 |
| 141Pr | Bcl-6 | 1:50 | EPR11410-43 | abcam | ab249707 | GR3296804-2 |
| 142Nd | PAX5 | 1:50 | D7H5X | CST | 12709BF | 2 |
| 143Nd | CD206 | 1:50 | E2L9N | CST | 91992BF | 2 |
| 144Nd | IDO | 1:50 | D5J4E | CST | 86630BF | 6 |
| 145Nd | CXCL13 | 1:50 | Polyclonal | Invitrogen | PA5-47035 | VI3068879 |
| 146Nd | PD-L1 | 1:25 | SP142 | abcam | ab236238 | GR3318567-1 |
| 147Sm | CD163 | 1:100 | EDHu-1 | Invitrogen | MA1-82342 | UL2904541 |
| 148Nd | CD14 | 1:50 | D7A2T | CST | 56082BF | 2 |
| 149Sm | CD11b | 1:50 | EPR1344 | abcam | ab209970 | GR3341282-3 |
| 150Nd | CD146 | 1:50 | EPR3208 | abcam | ab210072 | GR3356305-1 |
| 151Eu | CD31 | 1:50 | EPR3094 | abcam | ab207090 | GR3229164-6 |
| 152Sm | BCL2 | 1:100 | 124 | Novus Biological | NBP2-34513 | 596-3PABX200328 |
| 153Eu | CD44 | 1:50 | IM7 | BioLegend | 103001 | B223315 |
| 154Sm | Vimentin | 1:100 | D21H3 | CST | 5741BF | 9 |
| 155Gd | CD123 | 1:25 | IL3RA/1531 | abcam | ab268196 | GR3317070-4 |
| 156Gd | CD4 | 1:50 | EPR6855 | abcam | ab181724 | GR3285644-12 |
| 158Gd | PDPN | 1:50 | LpMab-12 | CST | 26981BF | 2 |
| 159Tb | CD68 | 1:100 | KP1 | Invitrogen | 14-0688-82 | 24B4431 |
| 160Gd | VISTA | 1:50 | D1L2G | SBT | 3160025D | 1351808 |
| 161Dy | CD20 | 1:100 | L26 | Invitrogen | 14-0202-82 | 2380076 |
| 162Dy | CD8a | 1:100 | C8/144B | Invitrogen | 14-0085-82 | 2151435 |
| 163Dy | CXCL12 | 1:25 | D8G6H | CST | 97958BF | 2 |
| 164Dy | Fibronectin | 1:50 | E5H6X | CST | 26836BF | 2 |
| 165Ho | HLA-DR | 1:100 | EPR3692 | abcam | ab215985 | GR3274622-7 |
| 166Er | CCL21 | 1:25 | Polyclonal | Invitrogen | PA5-84055 | VK3126119A |
| 167Er | CD209 | 1:50 | 8B6 | Invitrogen | MA5-15746 | VI3070941 |
| 168Er | Ki67 | 1:100 | B56 | BD | 556003 | 8239549 |
| 169Tm | CD49a | 1:50 | Polyclonal | Invitrogen | PA5-95563 | VI3070905A |
| 170Er | CD21 | 1:100 | SP186 | abcam | ab240987 | GR3350080-2 |
| 171Yb | SOX9 | 1:50 | D8G8H | CST | 82630BF | 2 |
| 172Yb | CD34 | 1:50 | EP373Y | abcam | ab198395 | GR3357088-1 |
| 173Yb | S100A9 | 1:400 | EPR3555 | abcam | ab227570 | GR3202573-1 |
| 174Yb | HLA Class I | 1:200 | EMR8-5 | abcam | ab70328 | GR3266685-2 |
| 175Lu | CD1a | 1:50 | EP3622 | abcam | ab217214 | GR3273995-1 |
| 176Yb | CD11c | 1:50 | EP1347Y | abcam | ab216655 | GR3357092-4 |

**Table S2.** Agent-based model parameters. Parameter values, justifications, and primary references for the agent-based model described in **Figure 6** and Methods. Organized by functional category: spatial geometry, initial cell counts, tumor B cell proliferation, CD8 T cell killing, suppression, chemokine fields, VISTA (contact-dependent), chemotaxis, immune turnover, and treatment interventions (anti-VISTA, rituximab, bispecific CD20-CD3). Parameters sourced from IMC-FL measurements, published literature, or calibrated so that the model reproduces key observed spatial phenotypes (immune exclusion gradient, Treg barrier, M2 Mac compartmentalization). The CD20-negative resistant fraction (3%) and CD20-low bispecific activity factor (0.5×) encode the clinically documented rituximab-escape mechanism and the higher avidity of modern bispecifics, respectively.

| Parameter | Value | Justification | Reference |
| --- | --- | --- | --- |
| *Spatial geometry* |  |  |  |
| Grid cell size | ~10 μm | Lymphocyte diameter 7-10 μm | BioNumbers BNID 100507 |
| FDC zone radius | 15 cells | FDC network inner 1/3 of follicle | Willard-Mack 2006 |
| B zone radius | 30 cells | FL follicle ~500 μm diameter | Willard-Mack 2006 |
| Boundary width | 5 cells | Treg enrichment at T-B border | Sayin 2018, JEM |
| Tissue radius | 45 cells | TMA core radius | This study |
| *Initial cell counts* |  |  |  |
| FDC | 40 | Rare stromal network | Park 2005, Immunology |
| Tumor B | 600 | Dominant (>75%) in follicle | FL histopathology |
| CD8 T | 120 | Yields ~2% follicular, ~28% T zone | This study, Fig 2 |
| CD4 T / Tfh | 80 | CD4:CD8 > 1; Tfh enriched | Pangault 2010, Leukemia 24:2080 |
| Treg | 60 | Treg:CD8 = 1.34 at boundary | This study, Fig 2 |
| M2 Mac | 80 | ~41% in FDC zone | This study, Fig 5 |
| M1 Mac | 50 | Interfollicular, pro-inflammatory | This study, Fig 5 |
| S100A9+ MDSC | 30 | ~3% of total; 3.6x in transformers | This study, Fig S11 |
| *Tumor B proliferation* |  |  |  |
| Proliferation rate | 0.005/tick | Ki-67 median ~16% in FL | Koster 2007, Haematologica 92:184 |
| Survival dependence | f(signal) | FDC niche required (BAFF, IL-15) | Laurent 2024, Blood 143:1080 |
| FDC survival emission | 0.5 | Primary anti-apoptotic source | Aguzzi 2014, Trends Immunol |
| *CD8 T cell killing* |  |  |  |
| Base kill rate | 0.08 | 2-16 kills/CTL/day in vivo | Halle 2016, Immunity |
| Activation bonus | 0.25 | Additive sublethal cytotoxicity | Weigelin 2021, Nat Commun |
| Exhaustion rate | 0.02 | Calibrated to 73.6% in GC | This study, Fig 2 |
| VISTA exhaustion | 0.008 | Contact-dependent quiescence | Johnston 2019, Nature |
| IDO exhaustion | 0.006 | Diffusible metabolite-driven | Liu 2018, Cancer Cell |
| *Suppression* |  |  |  |
| Background | 0.05 | Baseline immunosuppression in FL | This study, Fig 2 |
| Treg strength | 0.3 | CTLA-4, IL-10, TGF-β contact | Vignali 2008, Nat Rev Immunol |
| M2 Mac strength | 0.25 | Arginase, IL-10 suppression | Mantovani 2002, Trends Immunol; Lines 2014, Cancer Res |
| MDSC strength | 0.35 | Potent immunosuppressive activity | Veglia 2021, Nat Rev Immunol |

Table S2. Agent-Based Model Parameters (2/2)
| Parameter | Value | Justification | Reference |
| --- | --- | --- | --- |
| **Chemokine fields** |  |  |  |
| CXCL13 emission (FDC) | 0.4 | FDCs primary follicular source | Ansel 2000, Nature |
| CXCL13 decay | 0.01 | Slow (heparan sulfate binding) | Miller 2018, Front Immunol |
| CXCL9/10 emission (M1) | 0.05×act. | IFN-γ inducible | Groom 2011, Immunol Cell Biol |
| CXCL9/10 decay | 0.02 | Faster: inducible, transient | Groom 2011 |
| IDO emission (FDC zone) | 0.3 | Field-level; IMC FDC IDO partly M2 Mac spillover | Munn 2016, Trends Immunol |
| IDO emission (M2) | 0.2 | IFN-γ inducible in M2 | Munn 1999, J Exp Med |
| IDO emission (MDSC) | 0.25 | Strongest per-cell IDO | This study, Fig S11 |
| IDO decay | 0.08 | Fast: kynurenine t½ ~hours | Badawy 2017, Int J Tryp Res |
| **VISTA (contact-dependent)** |  |  |  |
| FDC vista level | 0.6 | CD14+ FDCs VISTA+ at protein | This study, Fig 5 |
| M2 Mac vista level | 0.5 | 56% myeloid VISTA+ | This study; Lines 2014, Cancer Res |
| MDSC vista level | 0.7 | Highest per-cell VISTA | This study, Fig S11 |
| **Chemotaxis** |  |  |  |
| CD8 → CXCL9/10 | p=1.0 | CXCR3-driven, highly motile | Groom 2011 |
| CD4 → CXCL13 | p=0.8 | CXCR5 follicle homing | Breitfeld 2000, J Exp Med |
| Treg → CXCL13 | p=0.7 | Moderate CXCR5; noise → boundary | Sayin 2018, JEM |
| M2 → CXCL13 | p=0.6 | Follicle-attracted | This study, Fig 5 |
| FDC / Tumor B | static | Sessile stromal / follicular cells | Allen 2008, Semin Immunol |
| **Immune turnover** |  |  |  |
| Death rate | 0.003/tick | T cell t½ 3-42 days (subpopulation-dep.) | Gossel 2017, eLife |
| Spawn probability | 0.15/deficit | Homeostatic replenishment | Bousso 2003, Nat Immunol |
| Spawn location | r = 0.7-1.0 R | Perivascular immigration | This study, Fig 2 |
| **Anti-VISTA** |  |  |  |
| VISTA block | 70% | Direct VISTA-PSGL1 blockade | Iadonato 2023, Front Immunol |
| IDO reduction | 30% | Indirect via VISTA signaling | Thisted 2024, Nat Commun |
| Kill bonus | +0.15 | Restored CD8 effector function | HMBD-002, ASCO 2021 |
| **Rituximab** |  |  |  |
| Kill rate | 0.012/tick | ADCC/CDC of CD20+ B cells | Cartron 2002, Blood |
| Penetration decay | 0.03 | Binding-site barrier | Fujimori 1990, J Nucl Med |
| CD20-neg fraction | 0.03 | Clinical CD20 loss under treatment | Czuczman 2008, Clin Cancer Res |
| CD20-neg prolif factor | 0.8 | Fitness cost of CD20 loss | Design: clinical PFS ~12-24 mo |
| **Bispecific CD20-CD3** |  |  |  |
| Kill bonus | +0.10 | T cell redirection (CR 60%) | Budde 2022, Lancet Oncol |
| Exhaustion bypass | 30% | Forced synapse overcomes TOX | Bacac 2018; Hutchings 2021 |
| Seek probability | 0.4/tick | Bispecific bridging redirects | Krupka 2016, Leukemia |
| CD20-low activity | 0.5× | Partial engagement (higher avidity) | Bacac 2018; Hutchings 2021 |

## Notes

https://doi.org/10.5281/zenodo.20612591

